# Chromosome condensation mechanically primes the nucleus for mitosis

**DOI:** 10.1101/2025.09.01.673438

**Authors:** Vanessa Nunes, Margarida Moura, Sara F. Silva, Débora Vareiro, Nicolas Auduge, Nicolas Borghi, Jorge G. Ferreira

**Affiliations:** Instituto de Investigação e Inovação em Saúde (i3S), Universidade do Porto, Porto, Portugal; Programa Doutoral em Biomedicina, Faculdade de Medicina do Porto, Universidade do Porto, Porto, Portugal; Université Paris-Cité, CNRS, Institut Jacques Monod, Paris, France; Dept. Biomedicina, Faculdade de Medicina do Porto, Universidade do Porto, Porto, Portugal

## Abstract

Accurate transition into mitosis driven by cyclin B1-CDK1 activity is essential to avoid chromosome segregation errors and preserve genome integrity. How this activity is spatially controlled to trigger mitotic onset remains unclear. Here, we show that chromosome condensation triggers an increase in nuclear envelope (NE) tension. This increased tension is required for nuclear translocation of cyclin B1 and dynein loading on nuclear pore complexes (NPCs), ensuring timely mitotic entry. Micromanipulation experiments further indicate this tension-dependent mechanism requires SUN proteins on the NE. Disruption of chromosome condensation leads to an accumulation of the G2 checkpoint kinase Wee1 and inhibition of CDK1 activity, which result in a temporary delay in mitotic entry. This delay can be overridden by increasing tension on the NE, which accelerates the nuclear translocation of cyclin B1 and dynein loading. We propose that mitotic onset is controlled by a chromosome-dependent NE tension mechanism that enables robust spatiotemporal coupling between chromosome condensation and the structural changes required for an efficient mitosis.

## Introduction

Mitosis is a highly regulated process, essential for organism development and homeostasis. A precise spatiotemporal control of mitosis contributes to the accurate partitioning of chromosomes that could, otherwise, lead to segregation errors and aneuploidy (Dantas *et al*, 2022; Furuno *et al*, 1999; Thompson & Compton, 2008). Underlying this spatiotemporal regulation is an intricate biochemical pathway, controlled by a complex formed by cyclin B1 and its associated cyclin dependent kinase (CDK) 1 (Lindqvist *et al*, 2009), that translocates to the nucleus (Gavet & Pines, 2010a) during the transition from G2 to mitosis (G2-M). Once inside the nucleus, this complex performs a series of well-characterized phosphorylations on key nuclear substrates that result in chromosome condensation (Abe *et al*, 2011), nuclear pore complex (NPC) dismantling (Linder *et al*, 2017) and nuclear lamina disassembly (Heald & McKeon, 1990), ultimately triggering nuclear envelope permeabilization (NEP) (Lindqvist *et al*., 2009) and irreversible mitotic commitment. Therefore, cyclin B1-CDK1 was proposed to coordinate the cytoplasmic and nuclear events required for timely mitotic entry (Gavet & Pines, 2010a; Gavet & Pines, 2010b). Nevertheless, the signal that triggers the initial nuclear accumulation of the complex, and therefore determines the timing of mitotic onset, remains elusive.

Disassembly of the nuclear envelope (NE) during the initial stages of cell division (Champion *et al*, 2017) gives rise to an “open mitosis”, characteristic of mammalian cells. This process requires the removal of NE membranes by dynein-mediated microtubule forces (Beaudouin *et al*, 2002; Salina *et al*, 2002). Dynein associates with the NE during early prophase by interacting with specific NPC components. Dynein can either bind Nup358/RanBP2 through its interaction with BicD2 (Splinter *et al*, 2010) or associate with Nup133 via its binding to CENP-F/NudE/EL (Bolhy *et al*, 2011). Importantly, during prophase, CENP-F is exported from the nucleus in a CDK1-dependent manner to associate with NPCs and enable dynein loading (Berto & Doye, 2018; Bolhy *et al*., 2011; Loftus *et al*, 2017). Therefore, during the G2-M transition, nuclear-cytoplasm transport must be tightly regulated to ensure the ordering of early mitotic events.

While the nucleus has been traditionally viewed merely as a structure that needs to be disassembled at mitotic entry, accumulating evidence reveals that multiple nuclear components play critical roles in regulating mitotic timing and fidelity. (Belaadi *et al*, 2022; Champion *et al*, 2019; Dantas *et al*., 2022; Lima *et al*, 2024; Turgay *et al*, 2014). Still, how these nuclear components mechanistically contribute to mitosis is poorly understood. Here, we investigated how the NE regulates mitotic onset. We show that mitotic chromosome condensation during prophase generates NE tension which, in turn, coordinates the cytoplasmic and nuclear events required for efficient mitosis.

## Results

### The nucleus is remodelled during mitotic entry

During the G2-M transition, the cell nucleus is extensively remodelled (Champion *et al*., 2017) to ensure an efficient mitotic spindle assembly (Lancaster *et al*, 2013). Whether these changes impact nuclear morphodynamics is not known. To address this, we used a combination of quantitative live-cell microscopy with micromanipulation in cells during late G2/prophase. These cells were readily identified by the presence of condensed chromosomes (Fig. 1A), one of the first signs of mitotic commitment (McIntosh, 2016). In a parallel approach, prophase chromosome condensation was further confirmed using fluorescence lifetime imaging (FLIM) of histone EGFP-H4, which revealed lower lifetime measurements when compared to interphase cells (Fig. EV1A), reflecting an increased chromosome compaction (Auduge *et al*, 2019). Then, to determine if mitotic chromosome condensation correlated with changes in the physical state of the nucleus, we measured NE morphodynamics in HeLa cells expressing POM121-3xGFP/H2B-RFP. These cells were seeded in line micropatterns to standardize nuclear shape and imaged with high temporal resolution (200 msec), using spinning disk microscopy (Castro *et al*, 2020) (Fig. 1A). Notably, prophase cells showed increased membrane displacements across the entire NE (Fig. 1B-E), when compared to their interphase counterparts. Moreover, most of these membrane movements were directed towards the nuclear interior (Fig. 1E, F), suggesting they might be dependent on inwards-directed active forces imposed on the NE (Introini *et al*, 2023; Schreiner *et al*, 2015). Next, we disrupted the microtubule (MT) network, since MT-generated forces were previously shown to trigger changes in NE dynamics in interphase cells (Hampoelz *et al*, 2011). Cells treated with nocodazole (Noco) presented a significant dampening of the overall NE movements (Fig. EV1B), but their inwards orientation was not significantly altered when compared to control cells treated with DMSO (Fig. EV1C). Together, these results indicate that the magnitude of NE movements partly depend on forces exerted by MTs, but their inwards orientation does not.

**Figure 1.**
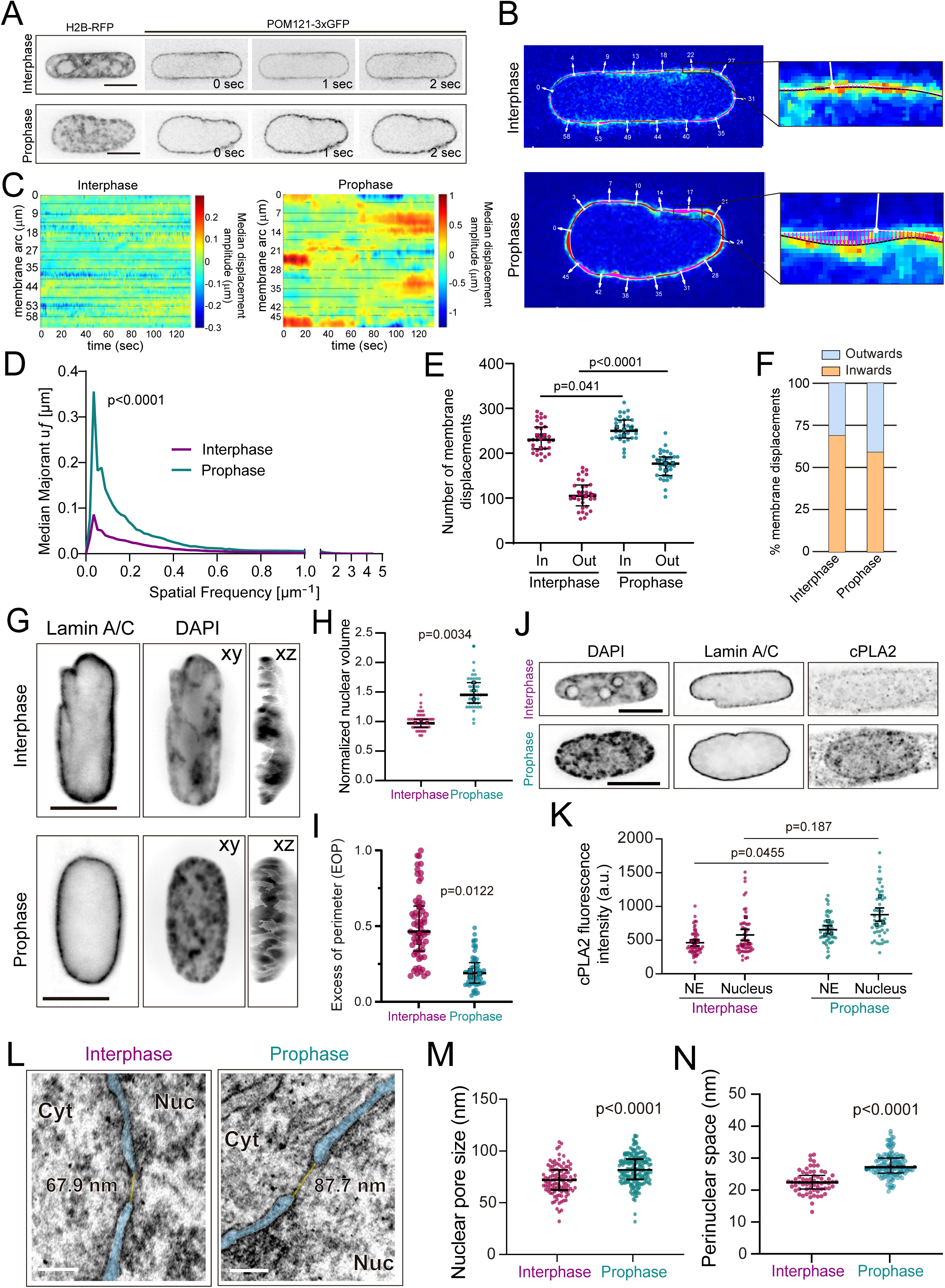
Nuclear remodelling during the G2-M transition. A) Selected frames from movies of HeLa cells expressing POM121-GFP/H2B-RFP, used for NE fluctuation analysis. Identification of prophase cells was done by assessing chromosome condensation with the H2B-RFP signal. B) Representative examples of inwards (pink vectors) or outwards (red vectors) NE membrane displacements of a cell in interphase (top) and a cell in prophase (bottom). C) Map of the membrane displacement amplitude, u, obtained for all time frames. The y axis corresponds to the arc around the median NE membrane. D) Median of the majorant of frequency-dependent displacements, *uf*, obtained by finding the maximum amplitude of the spatial Fourier transform (FT) for each frequency for interphase (n=32 cells from 4 independent replicates) and prophase cells (n=32 cells from 4 independent replicates; p<0.0001). This FT curve shows the maximum displacement amplitude of each wavelength. Quantification of the orientation (E) and proportion (F) of NE displacements for interphase and prophase cells (*p=0.041; p<0.0001). “Out” corresponds to outwards-directed movements and “In” corresponds to inwards-directed movements. G) Representative immunofluorescence images of nuclei labelled for Lamin A/C and stained with DAPI highlighting xy and xz projections. H) Quantification of nuclear volume for cells in interphase (n=51 cells, 3 replicates) and prophase (n=50 cells, 3 replicates; p=0.0034). I) Estimation of nuclear folding as assessed by the excess of perimeter (EOP) parameter for interphase and prophase cells (p=0.0122). J) Representative immunofluorescence images of nuclei labelled for cPLA2 and Lamin A/C and stained with DAPI. K) Quantification of cPLA2 fluorescence intensity on the NE and nucleus of interphase (n=55 cell, 3 replicates) and prophase cells (n=50 cells, 3 replicates; p=0.0455 for NE; p=0.187 for nucleus). L) Representative transmission electron microscopy (TEM) images of nuclear boundary regions in interphase and prophase. The NE is highlighted in blue and NPCs in yellow. Quantification of NPC size (M) and perinuclear space (N; PNS) for interphase (n=11 cells; 106 NPCs) and prophase cells (n=18 cells; 182 NPCs; p<0.0001).

Additional nuclear alterations were observed as cells prepared to divide. Our analysis revealed a significant increase in nuclear volume (Fig. 1G, H), which resulted in an unfolding of the NE, reflected by a decrease in the excess of perimeter (EOP) parameter (Fig. 1G, I). Both phenomena are indicative that, during the G2-M transition, the NE is under increased tension (Dantas *et al*., 2022). Accordingly, our analyses also revealed a recruitment of the cytosolic phospholipase A2 (cPLA2) to the NE (Fig. 1J, K), which normally occurs when the NE is under tension (Dantas *et al*., 2022; Enyedi *et al*, 2016; Lomakin *et al*, 2020; Venturini *et al*, 2020). Previous works have reported that imposing tension on the nucleus is sufficient to deform NPCs (Schuller *et al*, 2021; Zimmerli *et al*, 2021), leading to faster nuclear transport (Andreu *et al*, 2022; Dantas *et al*., 2022; Elosegui-Artola *et al*, 2017). Moreover, we have previously shown that nuclear tension during prophase can determine the timing of mitotic entry by regulating cyclin B1 translocation across the NPCs (Dantas *et al*., 2022). Therefore, we set out to confirm whether this increased nuclear volume and NE tension during prophase impacts NPC deformation. Comparison of NPC diameter between interphase cells and cells that were synchronized in G2 with a double thymidine block and subsequently released into prophase, indicated that synchronized cells present a significant dilation of NPCs (Fig. 1L, M; Fig. EV1E), which was accompanied by an increase in the perinuclear space (PNS; Fig. 1L, N). Together, these data demonstrate that a nuclear remodelling takes place during the G2-M transition.

### Chromosome condensation ensures the timing of mitotic entry

We then set out to determine the mechanism behind this prophase nuclear remodelling. We hypothesized the remodelling might be dependent on mitotic chromosome condensation, as this could be an early source of mechanical perturbation of the NE. To test this, we disrupted mitotic chromosome structure using well-established methodologies (Fig. 2A). We interfered with topoisomerase II (TopoII) activity using ICRF-193 (Ishida *et al*, 1994), inhibited the activity of histone deacetylases (HDAC) using valproic acid (VPA; Fig. EV2A, B) or depleted condensins using RNAi (Schneider *et al*, 2022) (SMC2 RNAi) (Fig. EV2C). Efficiency of these treatments in disrupting mitotic chromosome structure was assessed by analysing the coefficient of variation (CV) of chromatin (Neguembor *et al*, 2021). Notably, the treatments resulted in a significant decrease in the CV for the drugs (Fig. EV2D), as well as SMC2 RNAi (Fig. EV2E), indicating a disruption in chromosome condensation. FLIM measurements on EGFP-H4 confirmed the effect of ICRF-193, which abolished the decrease in fluorescence lifetime between interphase and prophase (Fig. EV2F). Although we cannot exclude additional side-effects of these treatments, it is unlikely that they would all lead to delays in mitotic entry independently of their role in chromosome condensation. Having established the efficacy of these treatments in disrupting mitotic chromosome condensation, we then asked whether they could lead to changes in NE morphodynamics. Treatment with ICRF-193 or VPA induced an increase in median membrane movements across the NE (Fig. EV2G), with a small but significant shift towards outwards-directed motions (Fig. 2B). These changes in NE dynamics indicate that chromosome condensation is likely responsible for the bias towards inward displacements during prophase, suggesting that condensation affects the tension state of the NE.

**Figure 2.**
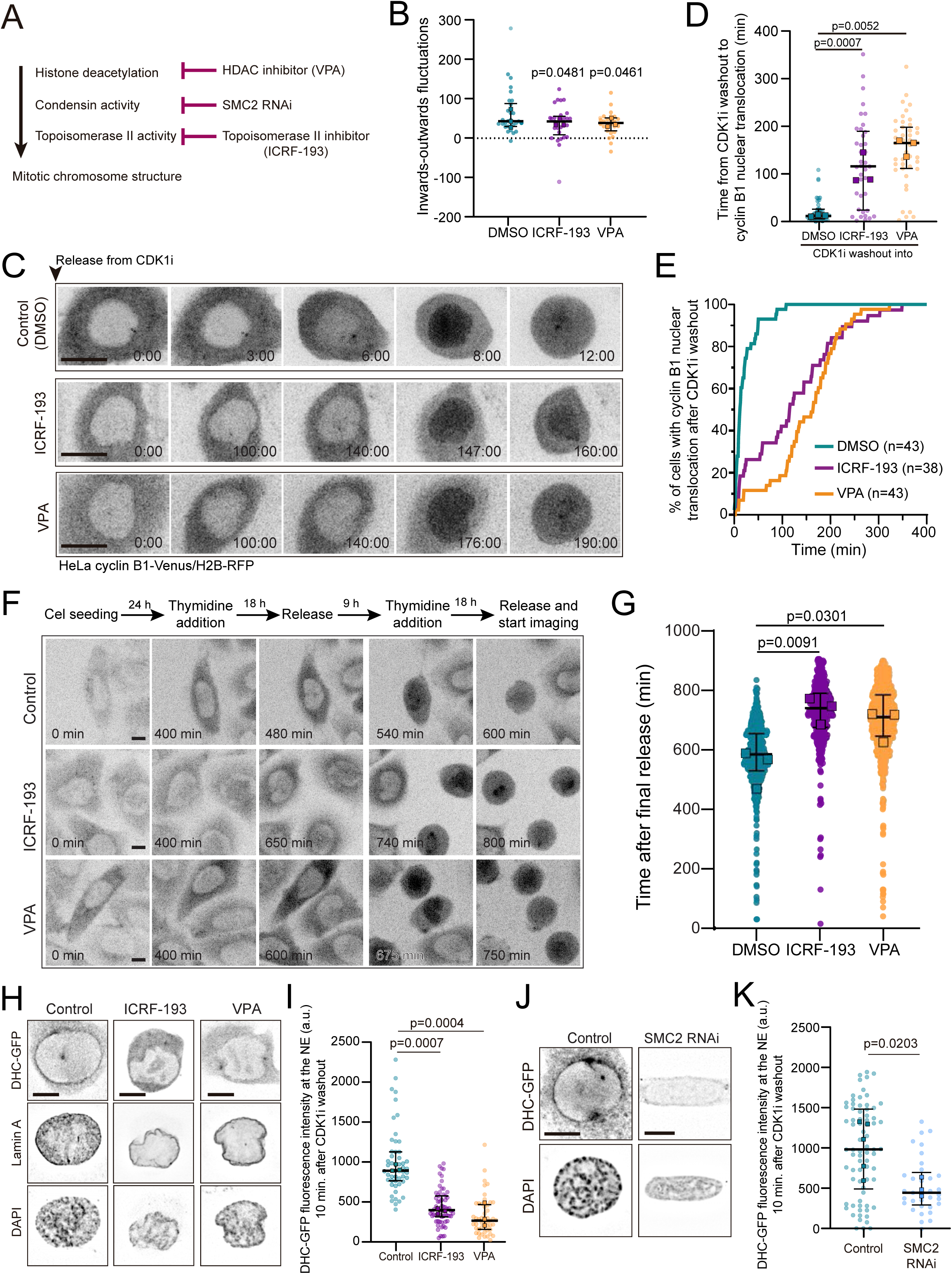
Chromosome condensation is required for timely mitotic entry. A) Scheme with the multiple strategies employed to interfere with mitotic chromosome structure. B) Quantification of the orientation bias in NE fluctuations for control cells (DMSO; n=30, 3 replicates) and cells treated with either ICRF-193 (n=29, 3 replicates; p=0.0481) or VPA (n=27, 3 replicates; p=0.0461). Values above zero reflect a bias towards inwards fluctuations. C) Representative frames from movies of cells expressing cyclin B1-Venus/H2B-RFP treated with DMSO (top panel), ICRF-193 (middle panel) or VPA (bottom panel). Time is represented in minutes (min) and zero corresponds to the moment of CDK1i washout. D) Quantification of the time for cyclin B1 nuclear translocation after CDK1i washout to for the indicated treatments (n=43 cells, 3 replicates for DMSO; n=38 cells, 3 replicates for ICRF-193, p=0.0007; n=43 cells, 3 replicates for VPA; p=0.0052). E) Cumulative percentage of cells with cyclin B1 nuclear translocation following CDK1i washout. F) Representative frames from movies of cells expressing cyclin B1-Venus after double thymidine synchronization treated with DMSO (top panel), ICRF-193 (middle panel) or VPA (bottom panel). Time lapse is 5 min. Time is in min. Scale bars, 10 μm. G) Quantification of the time from thymidine washout to cyclin B1 translocation for DMSO (n=680 cells, 3 replicates), ICRF-193 (n=356 cells, 3 replicates, p=0.0091) or VPA (n=902 cells, 3 replicates, p=0.0301). H) Representative immunofluorescence images of prophase cells expressing dynein heavy chain (DHC)-GFP after treatment with DMSO, ICRF-193 or VPA. I) Quantification of the levels of DHC-GFP fluorescence intensity on the NE for DMSO (n=52 cells, 3 replicates), ICRF-193 (n=63 cells, 3 replicates; p=0.0007) or VPA (n=49 cells, 3 replicates, p=0.0004) treated cells. J) Representative immunofluorescence images of DHC-GFP localization at the NE, following depletion of SMC2 by RNAi. K) Quantification of the levels of DHC-GFP on the NE following for control cells (n=74 cells, 5 replicates) or SMC2-depleted cells (SMC2 RNAi; n=33 cells, 3 replicates; p=0.0203). Scale bars, 10 μm.

Next, we asked whether perturbations in chromosome condensation could have implications for the timing and efficiency of mitotic entry. In preparation for mitosis, cyclin B1 translocates through the NPCs into the nucleus (Fig. EV3A), a process that depends on importins (Takizawa *et al*, 1999) and nuclear tension (Dantas *et al*., 2022). In addition, dynein is loaded on the NE (Fig. EV3B-D) in a CDK1-dependent manner (Baffet *et al*, 2015). We started by monitoring the nuclear translocation of cyclin B1 (Gavet & Pines, 2010a), as this event is required for NPC dismantling (Laurell *et al*, 2011) eventually leading to disassembly of the nuclear lamina and NEP. HeLa cells expressing endogenously tagged cyclin B1-Venus/H2B-RFP were synchronized in late G2 using a CDK1 inhibitor (CDK1i; RO-3306). Then, following release from the inhibitor, translocation of cyclin B1 into the nucleus was monitored using high-resolution live-cell imaging, as cells progressed towards mitosis (Fig. 2C; Movie S1). Notably, disruption of chromosome structure using ICRF-193 or VPA induced a significant delay in cyclin B1 nuclear transport (Fig. 2C-E; Fig. EV3E; Movie S1), with an increase in the half-time of translocation from 4.0±4.3 min in DMSO to 42.2±27.2 min in ICRF-193- (p=0.0036) and 67.4±49.6 min in VPA-treated cells (p=0.0001; Fig. EV3F). To rule out that cyclin B1 translocation delays could be due to the use of HeLa cells which have a perturbed cell cycle, we decided to use near-diploid, untransformed RPE-1 cells. Cells were treated with VPA, synchronized in late G2 with CDK1i and imaged during mitotic entry (Fig. EV3G). Importantly, treatment with VPA induced a comparable delay in cyclin B1 translocation (Fig. EV3H), confirming our earlier observations in HeLa cells. Although we cannot exclude other differences in cell cycle progression, our observations suggest that disrupting mitotic chromosome condensation is sufficient to delay cyclin B1 translocation in RPE-1 cells. Next, considering that prolonged inhibition of CDK1 could also impact the activity of other complexes such as cyclin A-CDK1, we decided to perform a double thymidine synchronization using HeLa cells expressing cyclin B1-Venus (Fig. 2F). Similarly to our previous results, treatment with ICRF-193 or VPA induced a significant delay in cyclin B1 translocation (Fig. 3G), independently of direct CDK1 inhibition. As a consequence of this delay in cyclin B1 translocation, DMSO-treated cells entered mitosis within 16.1±14.1 min after CDK1i washout, whereas both ICRF-193- and VPA-treated cells took on average 121.8±91.7 min and 150.5±72.3 min to enter mitosis, respectively.

**Figure 3.**
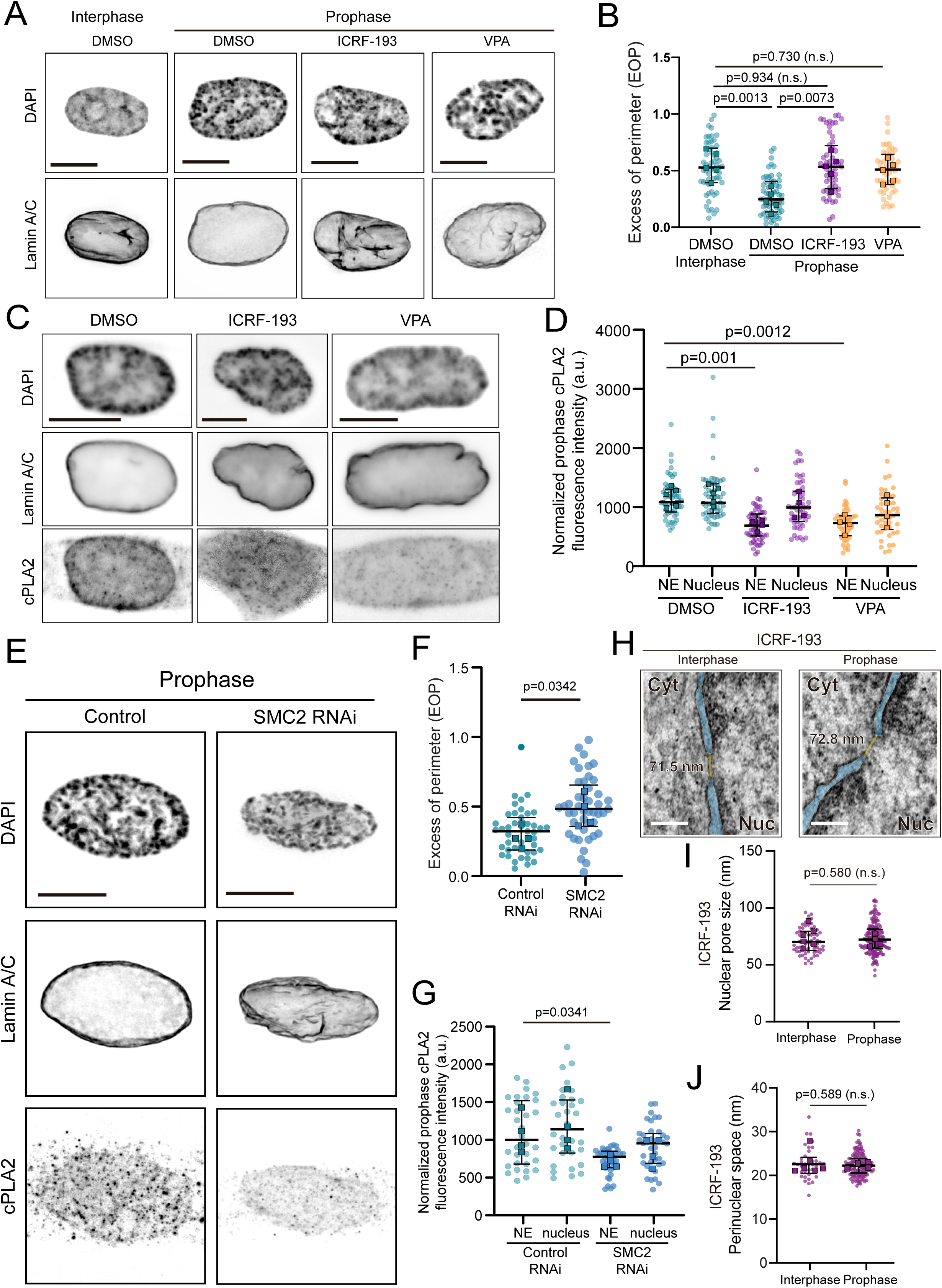
Nuclear envelope tension is determined by chromosome condensation. A) Representative immunofluorescence images of the NE of prophase cells following disruption of chromosome structure. B) Quantification of the excess of perimeter (EOP) in nuclei of interphase (n=59 cells, 5 replicates) and prophase (n=56 cells, 5 replicates) cells treated with DMSO, as well as prophase cells treated with ICRF-193 (n=56 cells, 5 replicates, p=0.0073) or VPA (n=49 cells, 3 replicates; p=0.0163). C) Representative immunofluorescence images of nuclei stained for cPLA2 in control cells (DMSO, n=53 cells, 5 replicates), and cells treated with ICRF-193 (n=52 cells, 4 replicates) or VPA (n=46 cells, 4 replicates) in prophase. D) Quantification of cPLA2 levels in the NE and nucleus of DMSO, ICRF-193 (p=0.001) or VPA (p=0.0012) treated prophase cells. E) Representative immunofluorescence images of nuclei stained for cPLA2 in control (scramble RNAi) or SMC2 depleted cells (SMC2 RNAi). F) Quantification of the EOP in nuclei of prophase cells treated with control (n=43 cells, 4 replicates) or SMC2 RNAi (n=45 cells, 4 replicates; p=0.0342). G) Quantification cPLA2 in the NE (p=0.0341) and nucleus of cells depleted of SMC2 by RNAi. H) Representative TEM images of the NE of cells treated with ICRF-193 in interphase (left panel) and prophase (right panel), highlighting the NE in blue and NPCs in yellow. Quantification of the nuclear pore size (I) and perinuclear space (J) in interphase (n= 6 cells; 67 NPCs) and prophase cells treated with ICRF-193 (n=17 cells; 168 NPCs). Scale bars in immunofluorescence images, 10 μm. Scale bars in TEM images, 100 nm.

To further confirm the contribution of chromosome condensation for mitotic entry, we then evaluated the dynamics of dynein loading on the NE. This process occurs by the binding of dynein to NPCs (Bolhy *et al*., 2011; Splinter *et al*., 2010), required for efficient NEP (Beaudouin *et al*., 2002; Salina *et al*., 2002) and early mitotic spindle assembly (Lima *et al*., 2024; Nunes *et al*, 2020). HeLa cells expressing dynein heavy chain (DHC) tagged with GFP were treated with either DMSO or ICRF-193 overnight and imaged during the G2-M transition (Fig. EV3L). In control prophase cells, dynein accumulated on centrosomes and the NE (top kymograph, black and yellow arrows, respectively). Then, following NE disassembly, dynein relocalized to kinetochores and the cell cortex (Fig. EV3M, yellow and red arrowheads, respectively). Strikingly, after treatment with ICRF-193, dynein no longer localized to the NE (Fig EV3L, bottom kymograph, yellow arrowheads), although centrosome, kinetochore and cortex localization were unaffected (Fig. EV3L, N). These results suggest that localization of dynein on the NE is dependent on the condensation state of chromosomes. To validate these observations, we extended our analysis by evaluating the localization of DHC-GFP on the NE of prophase cells following ICRF-193 or VPA treatment (Fig. 2H, I), or depletion of SMC2 by RNAi (Fig. 2J, K). HeLa cells were first synchronized in late G2 with CDK1i, released into prophase and fixed 10 min after release. As can be observed, interfering with mitotic chromosome structure leads to a significant decrease in the localization of dynein on the NE, when compared to control cells (Fig. 2H-K). Overall, these results demonstrate that chromosome condensation regulates the timing of key events required for mitotic entry.

### NE tension depends on chromosome condensation

We then set out to determine how chromosome condensation triggers the nuclear remodelling required for timely mitotic entry. During prophase, the nucleus unfolds and swells (Fig. 1G-I) leading to increased NE tension (Dantas *et al*., 2022). This increased tension regulates the dynamics of cyclin B1 nuclear translocation, setting the time for mitotic entry (Dantas *et al*., 2022). Based on this report and our current observations that cyclin B1 translocation (Fig. 2C-E) and NE dynein loading (Fig. 2H-K) are impaired upon disruption of chromosome structure, we hypothesized that NE tension could be dependent on efficient chromosome condensation. To test this, we treated cells with ICRF-193 or VPA and estimated the extent of NE folding by measuring the EOP parameter (Dantas *et al*., 2022; Lomakin *et al*., 2020). Here, increased values of EOP reflect a higher degree of NE folding. As can be observed, both treatments induced a significant increase in EOP of prophase nuclei, when compared to controls (Fig. 3A, B). As a result, these were now indistinguishable from nuclei of interphase cells (Fig. 3B). Next, we evaluated whether these treatments impacted NE tension. To assess this, we measured the nuclear levels of cPLA2, as these reflect increased tension on the NE in interphase (Enyedi *et al*., 2016; Lomakin *et al*., 2020; Venturini *et al*., 2020), as well as prophase cells (Dantas *et al*., 2022). Accordingly, both treatments induced a significant decrease in cPLA2 levels in prophase (Fig. 3C, D), when compared to control cells. These results were further corroborated in cells treated with SMC2 RNAi. Indeed, depleting condensin significantly increased the EOP (Fig. 3E, F), with a corresponding decrease in the levels of cPLA2 (Fig. 3G). Taken together, these results indicate that interfering with mitotic chromosome condensation impairs nuclear unfolding and decreases NE tension during prophase. The question then remains of how these changes might impact translocation of cyclin B1 across the NPCs. Recent works have shown that NPCs can deform *in vivo* and are sensitive to tension imposed on the NE (Schuller *et al*., 2021; Zimmerli *et al*., 2021), likely leading to increased nuclear transport (Elosegui-Artola *et al*., 2017). Moreover, we observed that prophase nuclei have an increased NPC diameter (Fig. 1J, K). These observations suggest that mitotic chromosome condensation increases tension on the NE, leading to NPC dilation and faster transport of cyclin B1 to the nucleus. To test this, we treated cells in either interphase or prophase with ICRF-193 and analysed NPC dilation and PNS size (Fig. 3H). After treatment, both NPC diameter and the PNS of prophase nuclei were undistinguishable from interphase nuclei (Fig. 3I, J), in contrast to what happens in untreated cells (Fig. 1J-L). Therefore, our results indicate that chromosome condensation impacts both NE structure and the timing of mitotic entry.

### Wee1 is required for the chromosome-dependent delay in mitotic entry

Next, we aimed to identify the targets downstream of chromosome condensation that could explain the delay in mitotic entry. For this, we analysed potential changes in expression levels and localization of key regulators of the G2-M transition in cells with disrupted chromosome condensation. Immunofluorescence analysis revealed that treatment with VPA or ICRF-193 increased the nuclear levels of Wee1 (Fig. 4A, B), an essential regulator of the G2-M transition (Heald *et al*, 1993; McGowan & Russell, 1993). Wee1 regulates mitotic entry by phosphorylating CDK1 on an inhibitory Y15 residue (CDK1 pY15), which blocks CDK1 activity (McGowan & Russell, 1993). Accordingly, immunofluorescence analysis using an antibody to specifically detect the levels CDK1 pY15 showed that disruption of chromosome condensation correlated with increased levels of this inhibitory phosphorylation (Fig. 4C, D). Overall, these results suggest that Wee1 sustains the delay in mitotic entry observed in cells with disrupted chromosome condensation. Accordingly, addition of the Wee1 inhibitor MK-1775 to VPA-treated cells during mitotic entry (Wee1i; Fig. 4E) was sufficient to allow the nuclear translocation of cyclin B1 within normal timings (Fig. 4F), abolishing the delay observed in cells treated with VPA only (Fig. 2). We did not observe changes in the levels of p21 (Fig. 4G, H) or p38 (Fig. 4I, J), suggesting that the observed delay does not depend on the p21-mediated DNA damage checkpoint (Bunz *et al*, 1998) or the p38-mediated antephase checkpoint (Matsusaka & Pines, 2004), respectively.

**Figure 4.**
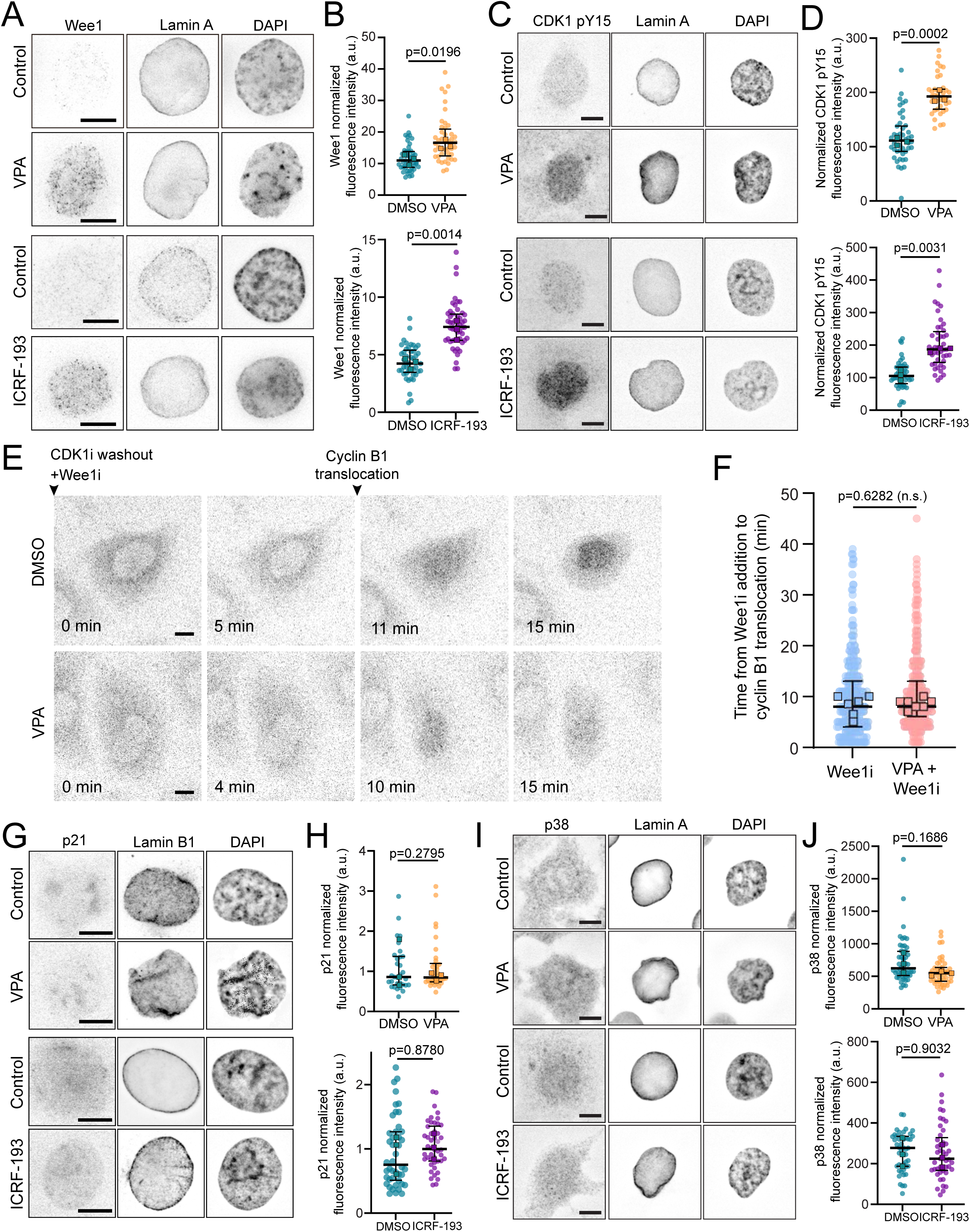
Wee1 delays mitotic entry when chromosome condensation is disrupted. Representative immunofluorescence images of cells stained for Wee1, Lamin A and DAPI for controls, ICRF-193 and VPA treated cells. B) Quantification of nuclear Wee1 levels in controls and VPA-treated cells (top plot; n=49 cells, 3 replicates for DMSO; n=44 cells, 3 replicates for VPA; p=0.0196). Quantification of nuclear Wee1 levels in controls and ICRF-193 treated cells (bottom plot; n=45 cells, 3 replicates for DMSO; n=52 cells 3 replicates for ICRF-193; p=0.0014). C) Representative immunofluorescence images of cells stained for CDK1 pY15, Lamin A and DAPI for controls, ICRF-193 and VPA treated cells. D) Quantification of CDK1 pY15 levels in controls and VPA-treated cells (top plot; n=46 cells, 3 replicates for DMSO; n=39 cells, 3 replicates for VPA; p=0.0002). Quantification of nuclear CDK1 pY15 levels in controls and ICRF-193 treated cells (bottom plot; n=45 cells, 3 replicates for DMSO; n=45 cells 3 replicates for ICRF-193; p=0.0031). E) Representative time frames of HeLa cells expressing cyclin B1-Venus, treated with DMSO or VPA. Cells were synchronized in late G2 using a CDK1 inhibitor (CDK1i, RO-3306). Then, cells were washed out of CDK1i and into medium with a Wee1 inhibitor (Wee1i, MK-1775) and filmed during mitotic entry. F) Quantification of the time from addition of the Wee1 inhibitor to the nuclear translocation of cyclin B1 for controls (Wee1i only; n=509 cells, 6 replicates) or VPA+Wee1i cells (n=485 cells, 7 replicates; p=0.6282). G) Representative immunofluorescence images of cells stained for p21, Lamin B1 and DAPI for controls, ICRF-193 and VPA treated cells. H) Quantification of p21 levels in controls and VPA-treated cells (top plot; n=31 cells, 3 replicates for DMSO; n=31 cells, 3 replicates for VPA, p=0.2795). Quantification of p21 levels in controls and ICRF-193 treated cells (bottom plot; n=44 cells, 3 replicates for DMSO, n=46 cells, 3 replicates for ICRF-193, p=0.8780). I) Representative immunofluorescence images of cells stained for p38, Lamin A and DAPI for controls, ICRF-193 and VPA treated cells. J) Quantification of p38 levels in controls and VPA-treated cells (top plot; n=50 cells, 3 replicates for DMSO, n=47 cells, 3 replicates for VPA, p=0.1686). Quantification of p38 levels in controls and ICRF-193 treated cells (bottom plot; n=46 cells, 3 replicates for DMSO, n=45 cells, 3 replicates for ICRF-193, p=0.9032).

### Imposing tension on the NE triggers mitotic entry

Since nuclear mechanics depends on the condensation state of chromatin (Stephens *et al*, 2017; Stephens *et al*, 2018) and its tethering to the NE (Schreiner *et al*., 2015), we now asked whether chromosome condensation regulates mitotic entry by increasing NE tension. To test this, we implemented two parallel approaches using either a cell confinement system or hypotonic treatment (Fig. 5A). Cell confinement induces NE unfolding and area expansion (Dantas *et al*., 2022; Lomakin *et al*., 2020), while hypotonic treatment results in an isotropic swelling of the nucleus and increased volume (Deviri & Safran, 2022). Using these approaches, we tested whether imposing tension on the NE of cells with compromised chromosome condensation was sufficient to restore normal mitotic entry. As a proxy for NE function in prophase, we used cells expressing cyclin B1-Venus to assess the dynamics of cyclin B1 translocation across NPCs, as well as cells expressing DHC-GFP to follow the dynamics of dynein accumulation on the NE, which is sensitive to the condensation state of chromosomes (Fig. 2H-K) and also required for efficient mitotic entry (Beaudouin *et al*., 2002; Nunes *et al*., 2020; Salina *et al*., 2002). In a previous work, we demonstrated that imposing tension on the NE induced a faster nuclear translocation of cyclin B1 (Dantas *et al*., 2022). Now, we tested whether artificially increasing NE tension could restore the dynamics of cyclin B1 translocation in cells with disrupted chromosome condensation. Indeed, cells that were treated with ICRF-193 or VPA and subjected to hypotonic shock quickly accumulated cyclin B1 in the nucleus (Fig. 5B, C). In addition, transiently increase NE tension in prophase cells treated with ICRF-193 was sufficient to restore dynein loading on the NE, similarly to control cells (Fig. 5D-G). To extend these observations, we next analysed NE dynein levels using an immunofluorescence approach in prophase cells that were previously treated with ICRF-193, VPA or SMC2 RNAi. Following treatment, confinement was applied to the cells for 30 min. After confinement release, cells were immediately fixed and processed. Notably, confinement was sufficient to promote NE dynein loading in ICRF-193- and VPA-treated cells (Fig. 5H, I) as well as in cells depleted of SMC2 (Fig. 5J, K), to levels similar to controls, overcoming the block induced by the treatments alone (Fig. 2H-K). These results indicate that increasing NE tension is sufficient to trigger mitotic entry, in conditions where chromosome condensation is disrupted.

**Figure 5.**
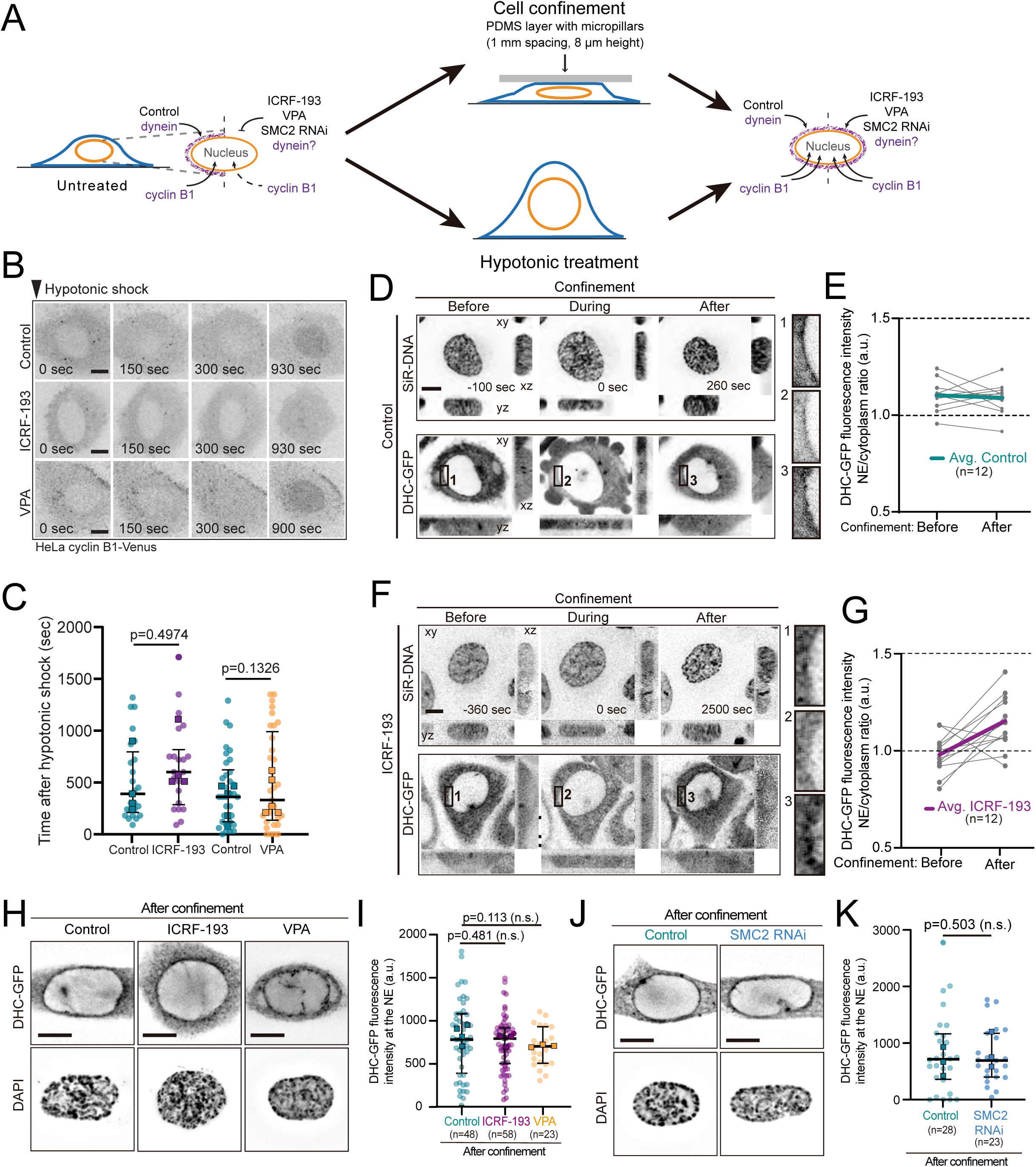
Increased NE tension imposed by chromosome condensation regulates mitotic entry. A) Schematic representation of the experimental setup designed to assess the response of the prophase nucleus to increased tension. B) Representative time frames from movies of cells expressing cyclin B1-Venus and treated with DMSO (top panels), ICRF-193 (middle panels) or VPA (bottom panels). Cells were subjected to hypotonic shock and filmed during mitotic entry. Time is in sec. Time lapse is 30 seconds. Scale bars, 10 μm. C) Quantification of the time from the hypotonic shock to cyclin B1 translocation for ICRF-193 (n=25 cells, 4 replicates for controls, n=22 cells, 4 replicates for ICRF-193, p=0.4974) or VPA (n=34 cells for controls, 5 replicates, n=33 cells for VPA, 5 replicates, p=0.1326). D) Representative images of control HeLa cell in prophase expressing DHC-GFP and stained with SiR-DNA under mechanical stimulation. Time zero corresponds to the moment when compression was applied to the nucleus. E) Ratio between the intensity of dynein on the NE and cytoplasm for control prophase cells (n=12). Note how the ratio is above 1, reflecting an enrichment of dynein on the NE during this stage of the cell cycle. F) Representative images of a prophase cell treated with ICRF-193 expressing DHC-GFP and stained with SiR-DNA, under mechanical stimulation. Time zero corresponds to the moment when compression was applied to the nucleus. G) Ratio between the intensity of dynein on the NE and cytoplasm for prophase cells treated with ICRF-193 (n=12). Note how the ratio increases following mechanical stimulation. H) Representative immunofluorescence images of DMSO, ICRF-193 or VPA treated cells after confinement, showing localization dynein on the NE. I) Quantification of DHC-GFP levels on the NE after mechanical stimulation (n=48 cells, 4 replicates for controls; n=58 cells, 3 replicates for ICRF-193, p=0.481; n=23 cells, 3 replicates for VPA, p=0.113). Note how ICRF-193 and VPA-treated cells accumulate dynein on the NE, similar to controls. J) Representative immunofluorescence images of controls (Scramble RNAi) and SMC2 depleted cells (SMC2 RNAi) showing dynein localization on the NE. K) Quantification of DHC-GFP levels on the NE of control cells (n=28 cells, 3 replicates) and cells depleted of SMC2 (n=23 cells, 3 replicates for SMC2 RNAi, p=0.503) after mechanical stimulation. Scale bars, 10 μm.

### The NE senses chromosome condensation through SUN proteins

To further dissect the mechanistic link between chromosome condensation, NE tension and mitotic entry, we systematically evaluated how different mechanosensitive NE components known to associate with chromatin, contribute to dynein loading and cyclin B1 translocation (Fig. 6A). Firstly, we disrupted the nuclear lamina in HeLa cells by depleting Lamin A using RNAi (Fig. EV4A). Decreasing Lamin A levels significantly changed the shape of prophase nuclei (Fig. 6B), which became irregular and deformed (Fig. 6C; Fig. EV4B), as determined by the nuclear irregularity index (NII), and showed an increase in membrane displacements across the entire NE (Fig. EV4C, D). Importantly, depleting Lamin A had no effect on dynein loading on the NE (Fig. 6D, E). These data suggest that the nuclear lamina, while necessary for maintaining nuclear shape, is likely not involved in the mechanical remodelling of the nucleus required for mitotic entry. Next, we asked whether this mechanical remodelling involves the LINC complex. This complex is comprised of KASH domain proteins such as Nesprins, that localize to the outer nuclear membrane (ONM) and bind to the cytoskeleton, and SUN proteins on the inner nuclear membrane (INM) that associate with chromatin, creating a physical link between the cytoskeleton and chromatin (Lombardi *et al*, 2011). We started by expressing a mutant form of KASH tagged with mRFP (DN-KASH) that binds to SUN proteins on the INM and displaces endogenous Nesprins from the ONM. This disrupts the interactions of the LINC complex with the cytoskeleton and blocks force transmission across the NE (Lombardi *et al*., 2011). As a control, we expressed a version of KASH that cannot interact with SUN proteins (KASH-ΔL) and therefore, does not interfere with endogenous Nesprin localization. As expected, expression of DN-KASH was sufficient to displace endogenous Nesprin-2 from the NE, which did not occur when KASH-ΔL was expressed (Fig. EV4E). Importantly, Nesprin displacement significantly decreased the levels of cPLA2 in the nucleus as well as in the NE (Fig. EV4F, G; p=0.0174 for NE and p=0.0288 for nucleus), indicating that an intact LINC complex is required for the increase in NE tension that occurs during prophase (Fig. 1J, K). These results are also in line with our previous observations in prophase cells showing that the LINC complex-mediated NE tension is required for timely translocation of cyclin B1 across the NPCs (Dantas *et al*., 2022). However, expression of DN-KASH did not affect the loading of dynein on the NE (Fig. 6F, G). Taken together, these results indicate that, while Nesprin localization on the ONM is required for efficient generation of tension on the NE and cyclin B1 translocation, it is not involved in the regulation of dynein loading. Finally, we asked whether SUN proteins on the INM were required for the mechanical remodelling of the nucleus during mitotic entry. Upon depletion of SUN2, similarly to cells expressing DN-KASH, we observed a significant delay in cyclin B1 nuclear translocation (Fig. 6H, I). Next, we evaluated whether SUN proteins were required for dynein accumulation on the NE. Depletion of either SUN1 or SUN2 resulted in a significant decrease in the levels of the respective protein (Fig. EV5A-C). This was accompanied by a decrease in dynein accumulation on the NE (Fig. 6J-L, EV5D), which correlated with decreased nuclear levels of cPLA2 (Fig. EV4H, I). Importantly, this loss of dynein in SUN-depleted cells occurred while the levels of chromosome condensation, as determined by measuring the CV, were even slightly increased, when compared to controls (Fig. EV5E). Overall, these data indicate that SUN proteins transmit the signal from mitotic chromosomes to the NE, ensuring timely cyclin B1 translocation and dynein loading, required for mitotic entry. Finally, we wanted to determine if increasing NE tension in cells depleted of SUN1 or SUN2 was sufficient to restore the timing of mitotic entry. Strikingly, confinement of SUN1 or SUN2 depleted cells was unable to rescue dynein loading on the NE (Fig. 6J, K). Similar results were obtained when both proteins were depleted simultaneously using a shRNA approach (Fig. EV5F), indicating that SUN1 and SUN2 likely play redundant roles in this process. Moreover, subjecting SUN2-depleted cells to a hypotonic treatment also failed to rescue dynein loading on the NE. (Fig 6L, M). Therefore, we concluded that SUN proteins are essential to transmit a mechanical signal from condensed mitotic chromosomes to the NE, to enable dynein localization and timely cyclin B1 translocation.

**Figure 6.**
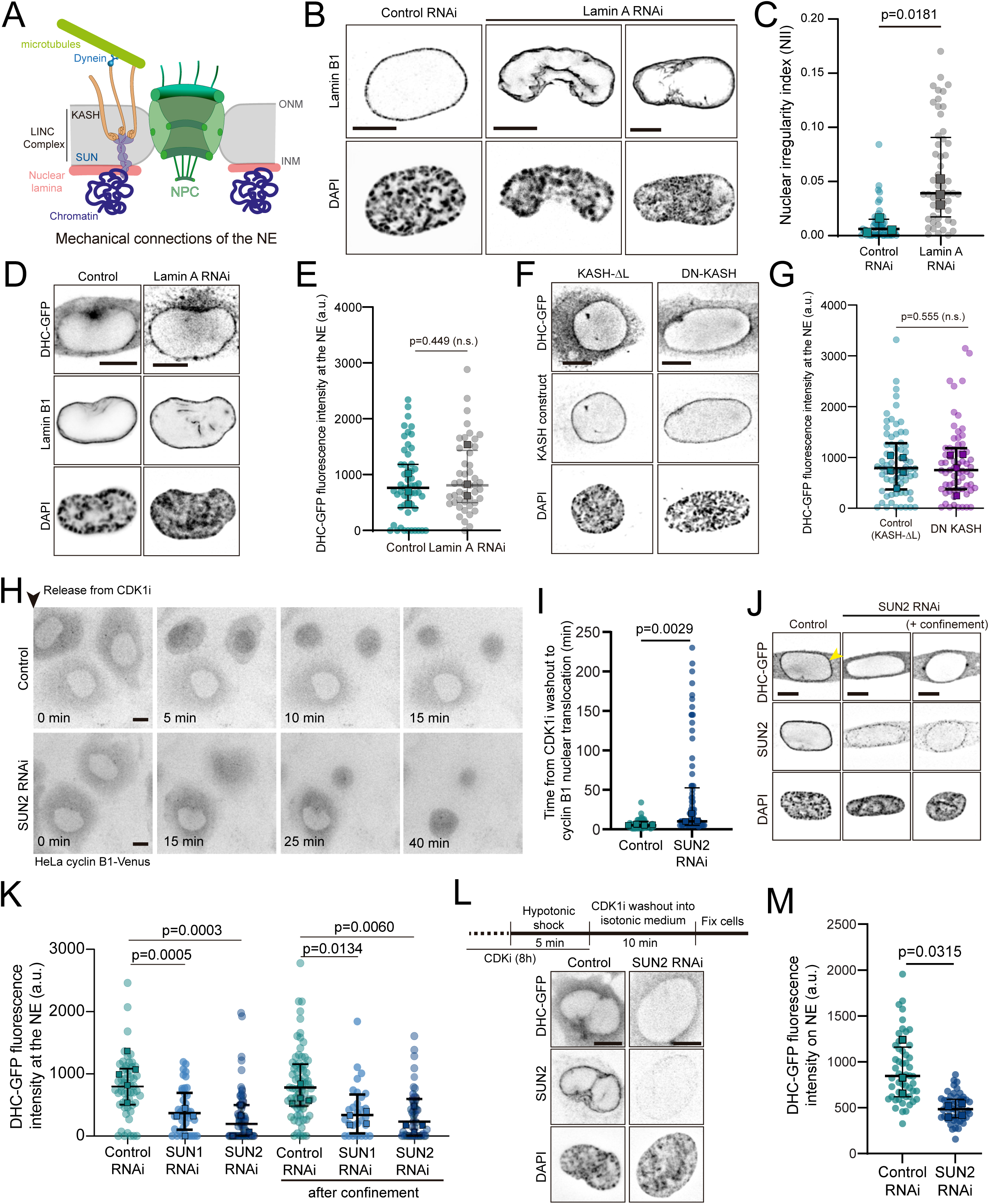
Chromosome condensation is coupled to NE through SUN proteins. A) Schematic depicting the multiple connections of chromatin to the NE. B) Representative immunofluorescence images of cells treated with scrambled RNAi (Control RNAi; n=51 cells, 3 replicates) or Lamin A RNAi (n=53 cells, 3 replicates; p=0.0181). C) Quantification of the nuclear irregularity index (NII) of control cells and Lamin A depleted cells (p<0.0001). This parameter was calculated as 1-solidity, with solidity being defined as nucleus area/nucleus convex area. D) Representative immunofluorescence images of a cell depleted of Lamin A, showing the localization of dynein on the NE. E) Quantification of dynein on the NE for control (n=53 cells, 3 replicates) or Lamin A depleted cells (n=49 cells, 3 replicates, p=0.449). F) Representative immunofluorescence images of cells expressing KASH-ΔL or DN-KASH, showing dynein localization on the NE. G) Quantification of dynein fluorescence intensity on the NE of cells expressing KASH-ΔL (n=68 cells, 5 replicates) or DN-KASH (n=65 cells, 4 replicates, p=0.555). H) Representative time frames of movies from control (top panels) and SUN2 RNAi (bottom panels) HeLa cells expressing cyclin B1-Venus. Cells were synchronized in late G2 with a CDK1 inhibitor. Following inhibitor washout, cells were filmed to monitor cyclin B1 nuclear translocation. Time lapse is 5 min. Time is in min. Scale bars, 10 μm. I) Quantification of the time from CDK1i washout to cyclin B1 nuclear translocation for controls (n=42 cells, 4 replicates) and SUN2 RNAi cells (n=105 cells, 3 replicates; p=0.0029). J) Representative immunofluorescence images showing dynein and SUN2 localization on the NE for controls and SUN2 depleted cells. K) Quantification of dynein levels on the NE for control cells (n=55 cells, 5 replicates), SUN1-depleted cells (n=42 cells, 4 replicates, p=0.0005) or SUN2-depleted cells (n=63 cells, 4 replicates, p=0.0003) without confinement. Similar quantifications were performed in controls (n=69 cells, 4 replicates), SUN1-depleted (=28 cells, 4 replicates, p=0.0134) or SUN2-depleted (n=45 cells, 4 replicates, p=0.0060) cells that were transiently confined (p<0.0001). L) Representative immunofluorescence images showing dynein and SUN2 localization. Cells were synchronized with a CDK1i and subjected to a hypotonic shock in the last 5 min. Then, cells were washed out of the inhibitor into isotonic medium. After 10 min, they were fixed and immunostained. M) Quantification of the levels of dynein on the NE for controls (n=53 cells, 3 replicates) and SUN2 RNAi cells (n=50 cells, 4 replicates, p=0.0315). Scale bars, 10 μm.

### A chromosome-SUN-NPC axis regulates mitotic entry

Based on our results showing that chromosome condensation affects the morphodynamics and tensional state of the NE in prophase, together with our previous work showing that NE tension promotes cyclin B1 nuclear translocation to control mitotic entry (Dantas *et al*., 2022), we now propose the following integrative working model: chromosome condensation drives an increase in nuclear volume and NE tension at prophase onset, leading to cPLA2 activation and opening of NPCs. In turn, this triggers a faster nuclear import of cyclin B1 and subsequent nuclear export of CENP-F to the ONM. Together, these translocations would allow the coordination of mitotic entry with chromosome condensation. To test this working model, we designed specific experiments that challenge some of its main predictions. We started by determining whether the loading of dynein on the NE, driven by mitotic chromosome condensation, was dependent on the nuclear export (Loftus *et al*., 2017) and subsequent accumulation of CENP-F on NPCs (Bolhy *et al*., 2011). Indeed, treatment with ICRF-193 or VPA led to a significant reduction of the levels of CENP-F specifically on the NE (Fig. 7A, B; yellow arrowheads; Fig. EV5G), while kinetochore binding was not affected (Fig. 7A, red arrowheads). A similar reduction was observed when we analysed the NE levels of NudE/EL, another component of the same dynein loading pathway (Fig. EV5H, I). Inversely, the levels of BicD2, involved in the alternative Nup358-BicD2 dynein loading pathway (Splinter *et al*., 2010) that does not require nuclear export, were not affected by ICRF-193 treatment (Fig. 7C, D; yellow arrowheads), even though chromosome condensation was significantly disrupted (Fig. EV5G). Overall, these data suggest that chromosome condensation, which we showed drives NPC dilation (Fig. 3H, I) and faster translocation of cyclin B1 into the nucleus (Fig. 2C-G), also facilitates the nuclear export and binding of CENP-F to NPCs to allow timely dynein binding.

**Figure 7.**
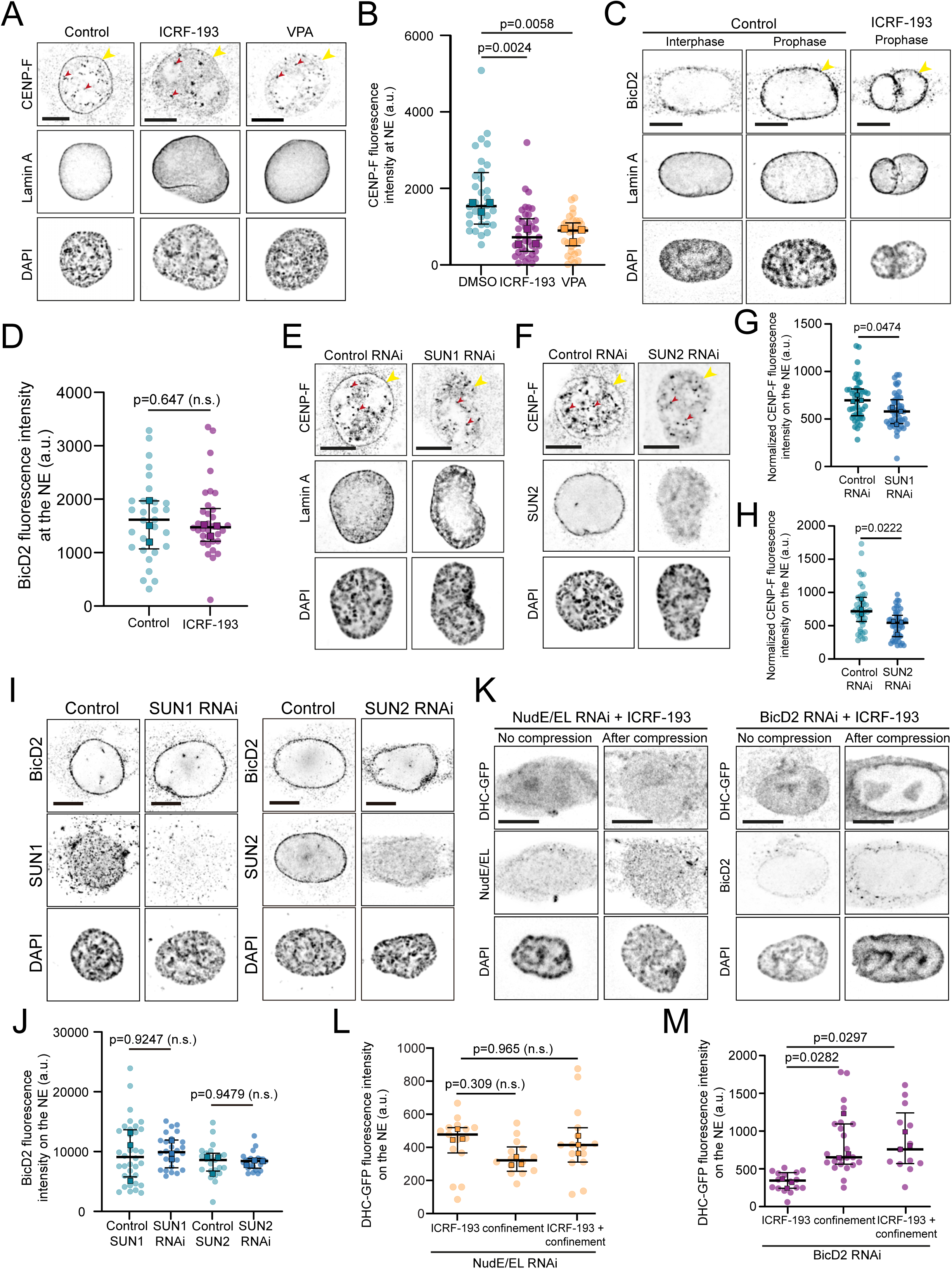
NE tension drives dynein accumulation through the Nup133-CENP-F/NudE/EL pathway. A) Representative immunofluorescence analysis of CENP-F localization in controls cells or cells treated with ICRF-193 or VPA. Yellow arrowheads indicate the NE; red arrowheads indicate kinetochores. B) Quantification of the levels of CENP-F on the NE for controls (n=32 cells, 3 replicates), ICRF-193 (n=39 cells, 3 replicates, p=0.0024) or VPA (n=36 cells, 3 replicates, p=0.0058). C) Representative immunofluorescence analysis of BicD2 localization in control cells (DMSO) or cells treated with ICRF-193. Note the accumulation of BicD2 on the NE of prophase cells (yellow arrowheads), irrespective of the treatment. D) Quantification of fluorescence intensity of BicD2 on the NE of control cells (n=29 cells, 3 replicates) or cells treated with ICRF-193 (n=34 cells, 3 replicates, p=0.647) during prophase. Representative immunofluorescence images of CENP-F localization in cells depleted of SUN1 (E) and SUN2 (F). Quantification of CENP-F fluorescence intensity on the NE of cells treated with SUN1 RNAi (G; n=45 cells, 3 replicates for controls and n=48 cells, 3 replicates for SUN1 RNAi, p=0.0474) or SUN2 RNAi cells (H; n=43 cells, 3 replicates for controls and n=49 cells, 3 replicates for SUN2 RNAi, p=0.0222), compared to their respective controls. I) Representative immunofluorescence images of BicD2 localization in control cells, as well as cells depleted of SUN1 (left panel) or SUN2 (right panel). J) Quantification of BicD2 fluorescence intensity on the NE following SUN1 (n=36 cells, 3 replicates for controls and n=26 cells, 3 replicates for SUN1 RNAi, p=0.9427) or SUN2 (n=23 cells, 3 replicates for controls and n=23 cells, 3 replicates for SUN2 RNAi, p=0.9479) depletion. K) Representative immunofluorescence images of dynein accumulation on the NE of prophase cells that were treated with ICRF-193 and depleted of NudE/EL (left panel) or depleted of BicD2 (right panel), without compression or after compression. L) Quantification of dynein fluorescence intensity on the NE following NudE/EL RNAi with either ICRF-193 treatment only (n=15 cells, 3 replicates), compression only (n=14 cells, 3 replicates, p=0.308) or ICRF-193 treatment with compression combined (n=15 cells, 3 replicates, p=0.965). M) Quantification of dynein fluorescence intensity on the NE following BicD2 RNAi with either ICRF-193 treatment only (n=16 cells, 3 replicates), compression only (n=23 cells, 3 replicates, p=0.0282) or ICRF-193 treatment with compression (n=13 cells, 3 replicates, p=0.0297). Scale bars, 10μm. N) Proposed model for the mechanical regulation of mitotic entry.

To further test our model, we then decided to disrupt NE tension in prophase cells (Fig. 1J, K) and assess whether this affected the localization of CENP-F on the NE. For this, we depleted SUN1 or SUN2, which are essential for transmitting the chromosome-mediated signal to the NE (Fig. 5J, K) and for ensuring NE tension during prophase (Fig. EV4H, I). Notably, depletion of either SUN1 or SUN2 resulted in a significant proportion of cells showing nucleoplasmic localization of CENP-F (Fig. EV5K), together with a reduction in the levels of CENP-F on the NE (Fig. 7E-H). On the other hand, no changes were observed in the levels of BicD2 on the NE when compared to controls (Fig. 7I, J). These data indicate that the increase in NE tension during prophase is required for the nuclear export of CENP-F, to allow timely loading of dynein on the NE through the Nup133-CENP-F-NudE/EL pathway. To further confirm these observations, we mechanically stimulated nuclei of cells that were treated with ICRF-193 and depleted of NudE/EL (Fig. EV5L). If this pathway is essential to ensure dynein loading, mechanical stimulation should not restore dynein on the NE. This would contrast with our previous results, where we could rescue dynein loading by mechanically stimulating nuclei that had compromised chromosome condensation, but otherwise normal levels of NudE/EL (Fig. 5H-K). Indeed, cells with disrupted chromosome condensation that were depleted of NudE/NudEL, could not localize dynein to the NE even after mechanical stimulation, making them undistinguishable from cells treated with ICRF-193 only (Fig. 7K, L). Inversely, mechanical stimulation of cells treated with ICRF-193 and depleted of BicD2 (Fig. S5L) was sufficient to rescue NE dynein loading (Fig. 7K, M). Taken together, these data indicate that the chromosome-mediated signal relayed by SUN proteins to the NE specifically targets the Nup133-CENP-F-NudE/EL pathway to ensure dynein loading and timely mitotic entry.

## Discussion

Mitotic entry is characterized by an extensive reorganization of cytoplasmic and nuclear structures (Champion *et al*., 2017; Gavet & Pines, 2010b), necessary for robust spindle assembly and chromosome segregation (Dantas *et al*., 2022; Lancaster *et al*., 2013). How these nuclear and cytoplasmic reorganizations are spatially and temporally coordinated to ensure an efficient mitosis remained unclear. Here we identify mitotic chromosome condensation as a key regulator of mitotic entry. By regulating NE tension, chromosome condensation controls the timing of cyclin B1 nuclear translocation and dynein association with NPCs, thereby determining irreversible mitotic commitment and ensuring an error-free mitosis (Dantas *et al*., 2022; Furuno *et al*., 1999).

In higher eukaryotes, mitotic entry is regulated by two mitotic cyclins, cyclin A and cyclin B. During the G2 phase, cyclin A localizes to the nucleus (Pines & Hunter, 1991) and controls the timing of mitosis (Furuno *et al*., 1999) by triggering the feedback loops required for initial CDK1 activation (De Boer *et al*, 2008; Hegarat *et al*, 2020). On the other hand, during the G2 phase cyclin B1 is mostly cytoplasmic (Pines & Hunter, 1991). As cells prepare to enter mitosis, cyclin B1 translocates into the nucleus where it is maintained until NEP (Gavet & Pines, 2010a). This initial translocation depends on cyclin A (Gong *et al*, 2007) and is further stimulated by a spatial positive feedback loop (Santos *et al*, 2012) that depends on NE tension (Dantas *et al*., 2022) and possibly involves changes in the nuclear import machinery (Gavet & Pines, 2010a). Once in the nucleus, the cyclin B1-CDK1 complex then further promotes chromosome condensation (Abe *et al*., 2011), NE dynein loading (Baffet *et al*., 2015), NPC disintegration (Laurell *et al*., 2011) and nuclear lamina disassembly (Heald & McKeon, 1990). Here, we propose that the signal for mitotic commitment depends on chromosome condensation, which increases NE tension (Fig. 1) to accelerate the nuclear accumulation of cyclin B1 (Fig. 2), subsequently triggering the downstream events referred above. This would require that the process of chromosome condensation start before the accumulation of cyclin B1 in the nucleus. The question then arises of how chromosome condensation could be initiated? One possibility is that activity of cyclin A in the nucleus could start the condensation process (Gong & Ferrell, 2010), leading to an increase in NE tension that would stimulate the translocation of cyclin B1 (Lindqvist, 2010; Santos *et al*., 2012), resulting in further chromosome condensation (Abe *et al*., 2011). This mechanism implies that, during the G2-M transition, the NE acts as a mechanosensor, detecting forces produced by chromosomes and relaying this signal to the cell cycle machinery. In fact, our results demonstrate that disruption of chromosome condensation increases the levels of Wee1, along with a corresponding Wee1-dependent inhibition of CDK1 (Fig. 4), necessary to sustain a delay in mitotic entry. This is in agreement with our previous observations showing that disruption of NE tension is sufficient to increase the levels of CDK1 pY15 and delay mitotic onset (Dantas *et al*., 2022). Under these conditions, cells would only enter mitosis once sufficient cyclin B1-CDK1 translocates to the nucleus to overcome the inhibitory effects of Wee1 (Lindqvist *et al*., 2009), leading to its proteosomal degradation (Watanabe *et al*, 2004). Therefore, by increasing NE tension and stimulating the nuclear translocation of cyclin B1, we tip the balance towards activation of the cyclin B1-CDK1 complex, which then results in the downstream nuclear events required for mitosis.

During the G2-M transition, forces exerted on the nucleus change considerably (Lima & Ferreira, 2024). As a result, the NE experiences a significant increase in tension that helps regulate the timing of mitotic entry (Dantas *et al*., 2022). However, the source of this tension remained unknown. We now propose that during the G2-M transition, the NE senses the tension imposed by condensing chromosomes, eliciting a downstream global cellular response (Fig. 8). On one hand, these forces are transmitted to the cytoplasm through the LINC complex, leading to increased actomyosin contractility (Dantas *et al*., 2022), NPC dilation (Fig. 1J) and fast nuclear accumulation of cyclin B1. On the other hand, these chromosome-derived forces also act specifically on the INM through SUN proteins, to enable the nuclear export of CENP-F (Fig. 7A, B) and dynein loading on NPCs. Accordingly, artificially increasing NE tension by confining cells or by hypotonic treatment was sufficient to restore mitotic entry, in cases where chromosome condensation was impaired. Surprisingly, dynein loading on the NE was not restored when confinement was applied to SUN-depleted cells, unlike what happens with expression of DN-KASH. This led us to hypothesize that SUN-dependent dynein loading relies on the transmission of forces to discreet sites near the NPCs, and not on a global increase of NE tension. This SUN-dependent, localized tension could alter NPC structure in way that is not recapitulated by the confinement setup, possibly explaining the lack of rescue under these conditions. In support of this hypothesis, it has been shown that SUN proteins and Nups can interact directly (Jahed *et al*, 2016; Liu *et al*, 2007; Talamas & Hetzer, 2011). Moreover, SUN-dependent forces can position NPCs (Smith *et al*, 2022) and generate tension islands on the NE that are sufficient to induce NPC dilation (Morgan *et al*, 2025). Nevertheless, the molecular details of this SUN-NPC interaction during the G2-M transition remain to be determined. Intriguingly, our results also indicate the nuclear lamina is not involved in this process. Depleting Lamin A did not affect dynein loading (Fig. 6D), although it significantly changed nuclear shape (Fig. 6B) and increased NE displacements (Fig. EV4D). Overall, these data suggest that the loading of dynein in not dependent on lamin-mediated nuclear mechanics but rather support the hypothesis that it requires the generation of forces in discreet spots at NPCs. It is well known that Interactions between chromosomes and the nuclear lamina are important for nuclear structure and mechanics during interphase (Herve *et al*, 2025; Schreiner *et al*., 2015; Stephens *et al*., 2017). However, during the G2-M transition, the nuclear lamina is disassembled in a CDK1-dependent manner to facilitate mitotic entry (Champion *et al*., 2017; Heald & McKeon, 1990). Together with our data, these observations suggest that chromosome condensation, and not the nuclear lamina, is the main contributor for NE tension during this stage of the cell cycle. Further experiments will be required to determine the contribution of each component of this chromosome-SUN-NPC pathway for nuclear mechanoresponse.

**Figure 8.**
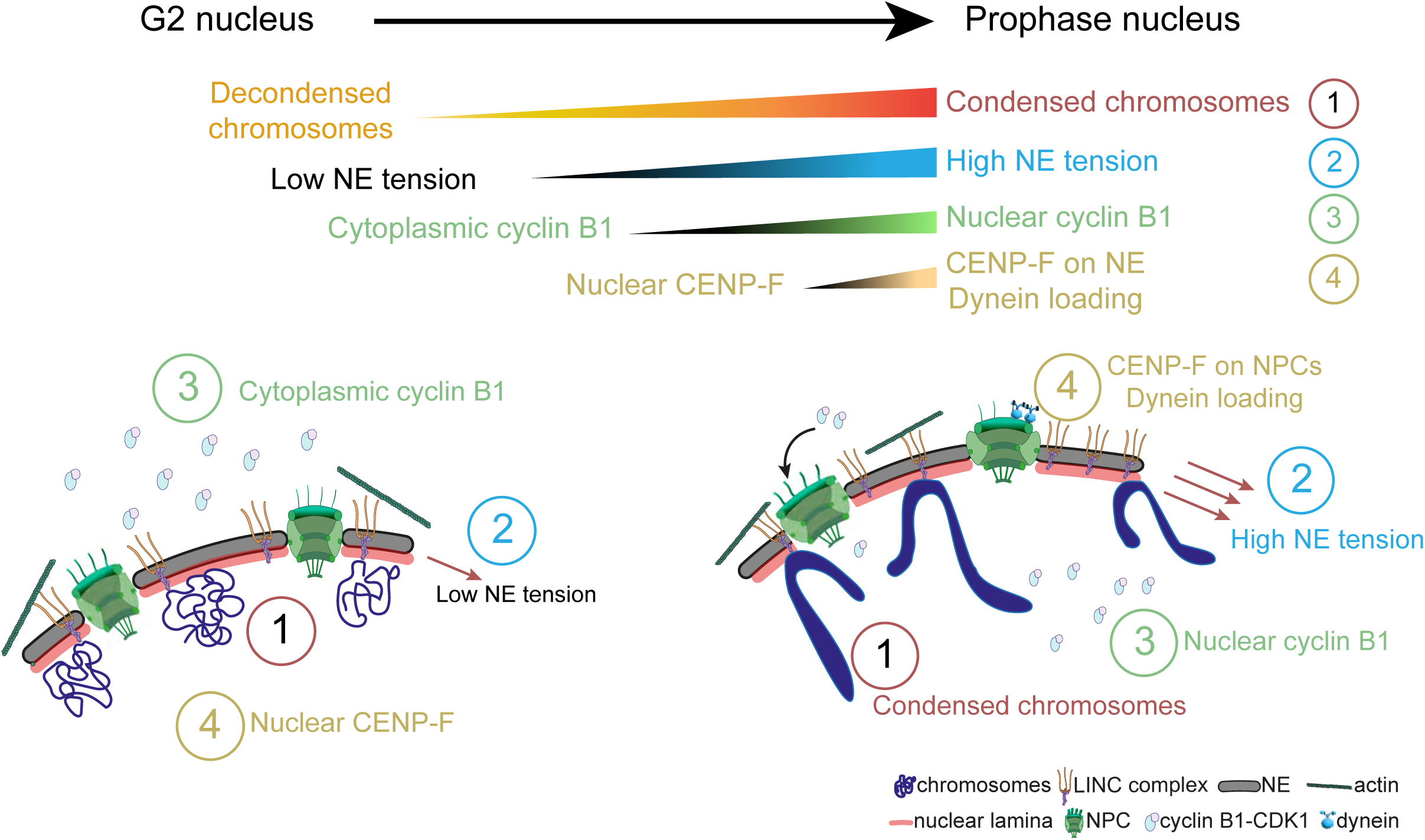
Proposed model for the regulation of mitotic entry by chromosome condensation. During the transition from G2 to mitosis, a series of coordinated events ensure the timing of mitotic onset. The first step involves mitotic chromosome condensation (1), which will lead to an increase in NE tension (2). As tension increases, NPCs dilate and cyclin B1 accumulates in the nucleus (3). Inside the nucleus, active cyclin B1-CDK1 will phosphorylate a series of nuclear targets, including CENP-F (4). This will lead to CENP-F export from the nucleus, enabling dynein loading on NPCs. Together, these allow cells to coordinate mitotic entry with chromosome condensation.

In eukaryotes, cyclins and their associated CDKs orchestrate the transitions between the different phases of the cell cycle (Basu *et al*, 2022). Deregulation of these transitions often lead to abnormal development (Kostic & Roy, 2002) and are associated with pathological conditions (Pellarin *et al*, 2025), as well as increased errors in chromosome segregation (Gayek & Ohi, 2016; Seibert *et al*, 2019). Importantly, recent works including our own, have demonstrated that cell cycle transitions are sensitive to mechanical stimuli (Aureille *et al*, 2019; Dantas *et al*., 2022; Donker *et al*, 2022; Gudipaty *et al*, 2017; Uroz *et al*, 2018). In this context, NE tension seems to be particularly relevant, as deformation of this organelle can regulate cell cycle progression by promoting the nuclear localization of key factors (Aureille *et al*., 2019; Dantas *et al*., 2022; Elosegui-Artola *et al*., 2017). Our results now indicate that timely transition into mitosis is dependent on the tension state of the NE that is regulated by chromosomes within the nucleus. Such a process would ensure that cells only commit to divide once chromosomes are sufficiently condensed, thus preventing the generation of errors caused by premature mitotic entry (Dantas *et al*., 2022; Furuno *et al*., 1999). We propose that mitotic entry is determined by the integration of biochemical and mechanical cues on the NE that enable a more robust coordination between the cytoplasmic and nuclear events, required for efficient cell division.

## Methods

### Cell lines

Cell lines were cultured in Dulbecco’s Modified Eagle Medium (DMEM; Life Technologies) supplemented with 10% fetal bovine serum (FBS; Life Technologies) and grown in a 37°C humidified incubator with 5% CO2. HeLa DHC-GFP line was a kind gift from Iain Cheeseman. HeLa CyclinB1-Venus cell line expressing endogenously tagged cyclin B1 was a gift from Jonathon Pines. HeLa POM121-3xGFP/H2B-mRFP cell line was kindly made available by Katharine Ullman. RPE-1 cell line expressing endogenously tagged cyclin B1-eYFP was kindly made available by Arne Lindqvist.

### Transient transfection and plasmids

For transient overexpression of mRFP1-KASH-DN or mRFP1-KASH-ΔL (kindly provided by Edgar Gomes), cells were transfected with the corresponding plasmid using Lipofectamine2000 (Invitrogen), according to the manufacturer’s instructions. Briefly, 5 μl of Lipofectamine 2000 and 0.6 μg/mL of DNA of interest were diluted and incubated in Opti-Minimal Essential Medium (Opti-MEM; ThermoFisher) for 30 min. Cells at 50%–70% confluence were then incubated with the DNA-lipid complexes for 6h. Prior to and during transfection, cells were cultured in reduced serum medium (DMEM supplemented with 5% FBS). Transfected cells were analyzed 72 h after transfection.

### RNAi experiments

Cells were transfected with small interfering RNAs (siRNAs) using Lipofectamine RNAi Max (Life Technologies), according to the manufacturer’s instructions. Briefly, 5 μl of Lipofectamine and each siRNA were diluted and incubated in Opti-MEM (ThermoFisher) for 30 min. Cells at 50%–70% confluence were then incubated with the siRNA-lipid complexes for 6h. Prior to and during transfection, cells were cultured in reduced serum medium (DMEM supplemented with 5% FBS). For all siRNAs used, cells were analyzed 48 - 72 h after transfection. Protein depletion efficiency was monitored by immunoblotting or immunohistochemistry. Different concentrations of siRNA were used: 20 nM for Lamin A RNAi, 40 nM for SUN1 and SUN2 RNAi and 200 nM for SMC2 RNAi. The following commercial ONTARGETplus siRNAs (Dharmacon) were used: Lamin A/C (#LQ-004978-00-0010), BicD2 (#L-014060-00-0020), SUN1 (#L-025277-00-0020), SUN2 (#L-009959-01-0020). For combined NudE/NudEL depletion the following oligos were ordered from Sigma-Aldrich 5′-GCUUGAAUCAGGCCAUCGA-3′ and 5′-UCGAUGGCCUGAUUCAAGC-3′ for NudE and 5′-GGAUGAAGCAAGAGAUUUA3′and 5′-UAAAUCUCUUGCUUCAUCC-3′for NudEL. For SMC2 depletion the following oligo was ordered from Sigma-Aldrich 5′-UGCUAUCACUGGCUUAAAUTT-3′. Both commercial ONTARGETplus non-targeting Pool siRNAs (Dharmacon #D-001810-10-20) and mock transfections were used as controls.

### Drug treatments

Inhibition of Topoisomerase II was done using 10 μM of ICRF-193 (Merck-Millipore). Inhibition of histone deacetylases (HDACs) was done using 1.5 μM of VPA (Sigma-Aldrich). Both drugs were added to the culture medium 8 – 16h before live-cell imaging or fixation. Control cells were treated with DMSO (Sigma-Aldrich) only.

### Cell synchronization

For the electron microscopy experiments cells were synchronized using a double-thymidine block. Cells were incubated with 2 mM thymidine (T1895; Sigma-Aldrich) for 16h, followed by a 10h release. To release the cells, the cells were washed three times in pre-warmed 1x PBS and incubated in pre-warmed fresh media. A second block was performed for another 16h, followed by an 8h release. To release the cells, the cells were washed three times in pre-warmed 1x PBS and incubated in pre-warmed fresh media containing either DMSO (D4540; Sigma-Aldrich) or ICRF-193 (Merck-Millipore).

For imaging experiments, cells were synchronized in late G2 using a CDK1 inhibitor (RO-3306; CDK1i; Sigma Aldrich SML0569). Cells were incubated with 10 μM RO-3306 for 10h-16h. Before imaging, the inhibitor was washed out three times using pre-warmed fresh medium and incubated in pre-warmed fresh media containing DMSO, ICRF-193 or VPA.

### Electron Microscopy

For ultrastructural analysis, the cells previously synchronized using a double thymidine block. Eight hours after the second release from the thymidine block, cells were fixed with 4% paraformaldehyde, 5% glutaraldehyde in 0.2M sodium cacodylate buffer for 15 min., with agitation. The fixative was removed and a new fixative solution of 2% paraformaldehyde, 2.5% glutaraldehyde, 0.1M sodium cacodylate was added to the cells for 1h. The samples were then post-fixed in 2% osmium tetroxide in 0.1 M sodium cacodylate buffer for 2h and then the cell pellet was resuspended in Histogel^TM^ and stained with a 1% aqueous uranyl acetate solution for 30 min, dehydrated and embedded in Embed-812 resin. Ultra-thin sections (60 nm thick) were sectioned on an RMC Ultramicrotome (PowerTome, USA) using a Diatome diamond knife, picked up on slot grids, and stained with uranyl and lead citrate for 5 min each. The samples were imaged on a JEOL JEM 1400 transmission electron microscope (JEOL, Tokyo, Japan) and the images were digitally recorded using an Orius 1100W CCD digital camera (Tokyo, Japan). Transmission electron microscopy (TEM) was carried out at the i3S HEMS center, Porto, Portugal. Nuclear pore complex (NPC) size was measured using Fiji(Schindelin *et al*, 2012).

### Micro-patterning

To control individual cell and nuclear shape, micro-patterns were produced as previously described (Azioune *et al*, 2009; Nunes *et al*., 2020). Briefly, glass coverslips were activated with plasma (Zepto Plasma System, Diener Electronic) for 1 min and then incubated with 0.1 mg/mL of PLL(20)-g[3,5]-PEG(2) (SuSoS) in 10 mM HEPES at pH 7.4, for 30 min, at room temperature (RT). After rinsing and air-drying, the coverslips were sealed onto a synthetic quartz photomask (Delta Mask), previously activated with deep-UV light (PSD-UV, Novascan Technologies) for 5 min, using 3 μL of MiliQ water. The coverslips were then irradiated through the photomask with the UV lamp for 5 min and incubated with 25 μg/mL of fibronectin (FBN; Sigma-Aldrich) in 100 mM NaHCO3 at pH 8.6, for 1 h, at RT. To monitor patterning efficiency and quality, 5 μg/mL of Alexa546 or 647-conjugated fibrinogen (Thermo Fisher Scientific) was added to the FBN solution. 50.000-75 000 cells were seeded per patterned-coverslip and allowed to adhere for 10–15 h before imaging. To wash out non-adherent cells, the cell medium was changed approximately 2–5 h after seeding.

### Cell confinement

For cell confinement experiments, confinement slides were fabricated using glass coverslips covered by a microstructured layer of PDMS, designed with a regular array of micropillars (diameter 449 μm, 1 mm spacing, 8 μm height), as described elsewhere(Le Berre *et al*, 2014). Briefly, a drop of polydimethylsiloxane (PDMS, RTV615, GE) mixture (8/1 w/w PDMS A/crosslinker B) was poured on top of a microfabricated mold. Glass coverslips with 10 mm diameter, previously activated with plasma for 2 min (Zepto system, Diener Electronics), were placed on top of the PDMS drop and gently pushed with tweezers to obtain a very thin layer of PDMS under the coverslips. The mold containing the coverslips was then baked on the hot plate at 95°C for 15 min. After removing the excess PDMS, the coverslips were released from the mold using isopropanol and a scalpel blade, rinsed with isopropanol and air dried. Prior to use, confinement coverslips were incubated for at least 1 h with the appropriate culture media and drugs (when necessary). The confinement slides were applied onto cells using either a dynamic or a static cell confiner. To perform dynamic cell confinement experiments(Le Berre *et al*., 2014), a custom-made suction cup made of a PDMS mixture (10/1 w/w PDMS A/crosslinker B) was baked on the hot plat at 80°C for 1h and left to dry overnight. After unmolding the device, a puncher (0.75 mm) was used to create a hole to plug the device into the vacuum generator apparatus (AF1-Dual, Elveflow). The confinement slide with the micropillars was then attached onto the piston of the PDMS suction cup and placed on top of a 35-mm imaging dish device. Cell confinement was modulated by increasing or decreasing the pressure on the vacuum line. For the static cell confinement experiments, we used a commercially available modified 6-well lid (4DCell). In this case, PDMS pillars were attached to the inside of the lid of the 6-well plate and confinement slides were attached to the bottom of the PDMS pillars. Once the lid was placed on the plate and locked, cells became confined. Confinement was removed by unlocking the lid from the plate.

### Time-lapse microscopy

For time-lapse microscopy, cells were seeded on FBN-patterned-coverslips one day before imaging. Before each experiment, the cell culture medium was replaced with Leibovitz’s-L15 medium (ThermoFisher Scientific) supplemented with 10% FBS and Antibiotic-Antimycotic solution (AAS; ThermoFisher Scientific). When SiR-dyes (20 nM SiR-tubulin or 10 nM SiR-DNA; Spirochrome) and/or drugs (acute pharmacological inhibition) were used, they were added to the culture medium before acquisition. Live-cell imaging was performed using temperature-controlled Nikon TE2000 microscopes equipped with a modified Yokogawa CSU-X1 spinning-disc head (Yokogawa Electric), an electron multiplying iXon+ DU-897 EM-CCD camera (Andor) and a filter-wheel. The following laser lines were used for excitation: 488, 561 and 647nm. The experiments were done with an oil-immersion 60x 1.4 NA Plan-Apo DIC (Nikon), except for the nuclear pore fluctuation analysis, in which an oil-immersion 100x 1.4 NA Plan-Apo DIC (Nikon) was used. Image acquisition was controlled by NIS Elements AR software. Approximately 17-21 z-stacks with a 0.5 μm or 0.7 μm separation were collected every 20 sec. For NE fluctuation analysis, a single z-stack was collected every 200 msec.

### Immunofluorescence

For CENP-F, cPLA2, SUN1, SUN2, Lamin A/C, Lamin B1, BicD2 and NudE/NudEL immunostainings cells were seeded on FBN-patterned-coverslips and fixed with 4% Paraformaldehyde (PFA) in Cytoskeleton Buffer (274 mM NaCl, 2.2mM Na2HPO4, 10mM KCL, 0.8 mM KH2PO4, 4 mM EDTA, 4 mM MgCl2, 10 mM PIPES, 10 nM Glucose, pH 6.1) for 10 min. Subsequently, cells were permeabilized with 0.5% Triton X-100 (Sigma- Aldrich) in 1x phosphate-buffered saline (PBS) for 5 min, and washed with 0.1% Triton X-100, 2x for 5 min. The cells were then blocked with 10% FBS in 0.1% Triton X-100 in PBS for 30 min. After blocking, the cells were incubated with the primary antibodies diluted in blocking solution, for 1 h at RT. Next, the cells were washed with 10% Triton X-100 in 1x PBS, 2x for 5 min and subsequently incubated with the respectively secondary antibody and DAPI (1ug/mL, Sigma-Aldrich), diluted in blocking solution, for 45 min at RT. Lastly, cells were washed with 0.1% Triton X-100 in 1x PBS, 2x for 5 min, and sealed on glass slides mounted with 20 mM Tris pH8, 0.5 N-propyl gallate and 90% glycerol. For SMC2 immunostainings cells were seeded on Poly-L-lysine-(PLL)-coated coverslips (50 μg/mL; Sigma, P8920). Cells were washed with 1x PBS for 1 min, pre-extracted with 0.5% Triton-X100 for 30 sec and fixed with 4% PFA in Cytoskeleton Buffer (274 mM NaCl, 2.2mM Na2HPO4, 10mM KCL, 0.8 mM KH2PO4, 4 mM EDTA, 4 mM MgCl2, 10 mM Pipes, 10 nM Glucose, pH 6.1) for 10 min. Next, cells were permeabilized with 0.5% Triton-X100 for 3 min, washed with 0.1% Triton-X100 for 2x for 3 min and blocked with 10% FBS in 0.1% Triton X-100 in PBS for 30 min. After blocking, the cells were incubated with the primary antibodies followed by the secondary antibodies and sealed on glass coverslips as described for the initial immunostainings. The following primary antibodies were used: mouse anti-Lamin A/C (1:500, Abcam), rabbit anti-Lamin B1 (1:500, Abcam), rat anti-alpha tubulin (1:500, Bio-Rad), rabbit anti-NudE/NudEL antibody (1:500, gift from Richard Vallee), rabbit anti-BicD2 (1:500, Atlas Antibodies) rabbit anti-SUN1 (1:1000, Merck), rabbit anti-SUN2 (1:1000, Merck), rabbit anti-SMC2 antibody (1:500; Bethyl Laboratories) rabbit anti-cPLA2 (1:100, #2832; Cell Signaling), rabbit anti-CENP-F (ab5, 1:300; Abcam). Alexa Fluor 488, 568 and 647 (1:2000, ThermoFisher Scientific) were used as secondary antibodies. Images were acquired using an AxioImager Z1 (63x, plan oil differential interference contract objective lens, 1.4 NA; all from Carl Zeiss) which is coupled with a CCD camera (ORCA-R2; Hamamatsu Photonics). Image acquisition was controlled by Zen software (Carl Zeiss).

### Quantification of fluorescence intensity on the nuclear envelope

Quantification of fluorescence intensity of NE proteins was performed using ImageJ, as described elsewhere(Lima *et al*., 2024). Briefly, a sum projection of three z-slices encompassing the central region of the nucleus was generated. On the sum-projected image, a segmented line (smoothened by a spline fit) of a defined width (w1) was drawn along the NE, and the transverse-averaged fluorescence signal (S1), containing the fluorescence signal as well as background, was measured. A second equivalent measurement (S2) was done using the same line, after increasing its width to w2. While the signal of interest remains the same in the dilated line, background increases by the factor w2/w1, which allows retrieval of I(r), the background-corrected profile, using the equation:

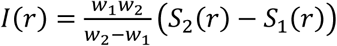

Line width w1 should be large enough to fully encompass the signal of interest, while w2 should be at least 20% larger than w1, while small enough to avoid inclusion of extraneous signal from non-NE sources. For our quantifications, the intensity profile (i.e. the r-dependence) was irrelevant, so I(r) was integrated along the full length of the curve and divided by the line length (or, equivalently, the line ’area’).

### CyclinB1 quantification

For quantifications of cyclin B1 levels and translocation rates, images were analysed using ImageJ. A small circular region of interest (ROI) was defined, and cyclin B1 fluorescence intensity measured, throughout time in the cell nucleus. The same ROI was used to measure the background outside the cell area. All fluorescence intensity values were then background corrected and area normalized. The values obtained were then fitted with the sigmoidal function 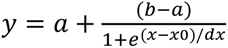 and scaled to their fitted maximum and minimum values. From this, we obtained cyclin B1 translocation rates and half-time. Time zero is defined as the moment when the CDK1 inhibitor is washed out.

### Quantification of nuclear volume, nuclear envelope excess of perimeter (EOP) and Coefficient of Variation (CV) of DNA

The excess of perimeter (EOP) parameter was calculated to estimate the amount of nuclear envelope (NE) area stored in folds, as described previously(Lomakin *et al*., 2020). To calculate the EOP, we measured the perimeter (P) and surface area (A) from 2D immunofluorescence images taken at the maximum radius of the nucleus, using Lamin A/C labelling. Next, we defined R0 as the radius of the circle defined by the area A. EOP was obtained as the ratio between (P - 2πR0) and (2πR0). Highly folded nuclei have EOP values close to 1, whereas in nuclei with smooth surfaces, EOP tends to zero.

The nuclear volume was estimated using ImageJ. Briefly, after image thresholding (Otsu method), the area of each slide in the stack was measured individually. The area measurements were then added together and multiplied by the depth of each slice. Finally, the nuclear volume measurements were normalized to the smallest and largest value in the data set using GraphPad.

Chromosome condensation was assessed usings the coefficient of variation (CV). This method evaluates the degree of heterogeneity of the DNA signal in the nucleus using DAPI staining(Neguembor *et al*., 2021), providing a simple way to quantify changes in overall chromatin compaction.

### Nuclear Envelope Dynamics

Nuclear envelope movements were quantified using the “Membrane fluctuations” package in Trackosome(Castro *et al*., 2020). Movements are defined as the distance from each point of the median membrane to the membrane at a given frame, along a direction normal to the median membrane. The membranes are segmented and centered for all frames, assuring a common centroid among frames. To obtain a reference membrane of the NE, we calculate the median projection of the centered frames and segment the resulting membrane. Each point of the reference membrane is then associated with a normal vector defining the direction of the membrane displacements. The movements are measured by calculating the distance between the reference membrane and the membrane at each frame, along the directions defined by the normal vectors.

### Fluorescence lifetime imaging (FLIM) of chromosome condensation

FLIM measurements to determine the degree of chromosome condensation are based on an approach described elsewhere(Auduge *et al*., 2019), with a different instrumentation as follows. Briefly, cells were imaged on a Zeiss LSM 980 (Zeiss, München, Germany) in combination with the LSM Upgrade Kit provided by PicoQuant (PicoQuant, Berlin, Germany). The LSM Upgrade Kit consists of a pulsed laser source at 480 nm generating nanosecond pulses at a repetition rate of 40MH injected into the LSM 980 and a PMA cooled hybrid photomultipliers plugged on a MultiHarp 150 Multichannel time correlated single photon counter which collects the fluorescence signal emitted from the sample through a spectral filter (BrightLine HC 520/35, Semrock). Images were analysed on SymPhoTime software (PicoQuant, Berlin, Germany). Fluorescence decay from chromatin-containing regions of interest is fitted by a monoexponential model, taking into account the instrumental response, yielding the fluorescence lifetime of the EGFP probe.

### Western Blotting

HeLa cell extracts were collected, washed, and resuspended in 30-50 μL of Lysis Buffer (20 nM HEPES/KOH pH 7.9; 1 mM EDTA pH 8; 1 mM EGTA; 150 mM NaCl; 0.5% NP40; 10% glycerol, 1:50 protease inhibitor (ROCHE); 1:100 Phenylmethylsulfonyl fluoride). The samples were then flash frozen in liquid nitrogen and kept on ice for 30 min. After centrifugation at 14000 rpm for 5 min at 4°C, the supernatant was collected, and protein concentration was determined by the Bradford protein assay (Bio-Rad). 50 μg of each protein were loaded and ran on 15% SDS-PAGE gels. Next, the loaded proteins were transferred to a nitrocellulose Hybond-C membrane using the iBlot Gel Transfer Device (Thermo Scientific). Afterwards the membranes were blocked with 5% Milk in triphosphate buffered saline (TBS) with 0.1% Tween-20 (TBS-T) for 1h at RT and subsequently incubated with the primary antibodies overnight at 4°C with shaking. After three washes in TBS-T, the membranes were incubated with the secondary antibody for 1h at RT. After several washes with TBS-T, the detection was performed with Clarity Western ECL Substrate (Bio-Rad). The following primary antibodies were used: mouse anti-Lamin A/C (1:500, Abcam), mouse anti-Nde1 (1:500, Abnova), rabbit anti-BicD2 (1:250, Atlas Antibodies) and rat anti-alpha tubulin (1:1000, Bio-Rad). The secondary antibodies used were anti-mouse-HRP, anti-rabbit-HRP and anti-rat-HRP at 1:5000.

### Statistical analysis and data presentation

Three to six independent experiments were used for statistical analysis. Knockdown efficiency was assessed by immunofluorescence or immunoblot quantification, as indicated. In all figures, and for visualization purposes only, fluorescence images were deconvolved using ImageJ and maximum intensity projections generated. Quantifications of fluorescence intensity were done using sum-projected images without any further manipulations. Plots show all data points, with the size of the bars (whiskers) ranging from the 25^th^ to the 75^th^ percentile and the line representing the median of the sample. Square dots represent the median of the replicates in each experiment. Statistical analysis for multiple group comparison was performed using a parametric one-way analysis of variance (ANOVA) when the samples had a normal distribution. Otherwise, multiple group comparison was done using a nonparametric ANOVA (Kruskal-Wallis). Multiple comparisons were analyzed using either post-hoc Student-Newman-Keuls (parametric) or Dunn’s (nonparametric) tests. When only two experimental groups were compared, we used either a parametric t test or a nonparametric Mann-Whitney test. Comparison for multiple time-course datasets was done using an ANOVA Repeated Measures, when the samples had a normal distribution. Otherwise, group comparison was done using Repeated Measures ANOVA on Ranks. Distribution normalities were assessed using the Kolmogorov–Smirnov test. No power calculations were used. All statistical analyses were performed using GraphPad Prism 9.5.1 (GraphPad Software).

## Resource availability

### Materials availability

Stable human cell lines generated in this study are available upon request.

### Data availability

The data supporting the findings of this study, as well as any additional information, are available from the corresponding author upon request. Source data are provided with this paper.

### Author contributions

V.N. performed experiments, analysed data, conceptualized the work and co-wrote the manuscript. M.M. performed experiments related with cyclin B1 translocation and CENP-F binding to the NE. S.V.S. performed experiments related cyclin B1 translocation and dynein loading following hypotonic shock in HeLa cells, as well as with cyclin B1 translocation in RPE-1 cells. D.V performed experiments related with HDAC inhibition and dynein accumulation on the NE. N.A and N.B performed the FLIM measurements and DNA condensation analysis. J.G.F. performed proof-of-concept experiments, analysed the data, provided funding, conceptualized the work and co-wrote the manuscript.

## Acknowledgments

The authors would like to thank Drs. Matthieu Piel and Edgar Gomes for critical reading and review of the manuscript. We thank Dr. Helder Maiato and the members of the CID lab for the helpful discussions and feedback. We further thank Dr. Helder Maiato for access to microscopy equipment. The authors thank Dr. Katharine S. Ullman for the HeLa POM121-3xGFP cell line, Dr. Richard Vallee for the NudE/EL antibody, Dr. Edgar Gomes for the KASH-ΔL and DN-KASH constructs, Dr. Joanthon Pines for the HeLa cyclin B1-Venus cell line and Dr. Arne Lindqvist for the RPE-1 cyclin B1-eYFP cell line. The authors would like to thank Sofia Pacheco and Rui Fernandes at the HEMS facility at i3S for assistance with the TEM analysis.

This work was supported by Fundação para a Ciência e Tecnologia IP, grant reference “ERC-PT A-Projects – MECHANIST”, through the measure RE-C06-i06 – “Ciência Mais Capacitação” of the Recovery and Resilience Program, attributed to J.G.F. D.V. is supported by a PhD fellowship PRT/BD/154964/2023, from Fundação para a Ciência e Tecnologia (FCT), I.P., in the scope of the European University Alliance for Global Health (EUGLOH) Consortium. This work was supported by the Fondation pour la Recherche Médicale, grant number EQU202203014613 to NB. NA and NB acknowledge the ImagoSeine facility, member of the France BioImaging infrastructure (ANR-10-INSB-04).

## Expanded View Figure Legends

**Figure EV1 – Microtubule dependent changes in NE membrane displacement**

A) Fluorescence lifetime microscopy (FLIM) measurement of histone H4-GFP of cells in interphase (n=50 cells, 3 replicates) and prophase (n=36 cells, 3 replicates, p<0.0001). Decreased fluorescence lifetime in prophase cells reflects increased chromosome compaction. B) Median of the majorant of frequency-dependent NE membrane displacements, uf, obtained by finding the maximum amplitude of the spatial Fourier transform (FT) for each frequency for DMSO (n=30, 3 replicates) and Nocodazole treated cells (Noco; n=27, 3 replicates; p<0.0001). This FT curve shows the maximum displacement amplitude of each wavelength. Quantification of the orientation (C) and proportion (D) of NE membrane displacements for DMSO and Noco cells (p=0.0023). E) Representative TEM images of interphase (left) and prophase (right) nuclei. Scale bars correspond to 5 μm.

**Figure EV2 – Disrupting chromosome condensation affects NE membrane displacements**

A) Representative immunofluorescence images of histone H3 acetylation on lysine 9 (H3K9ac) in controls (DMSO) and VPA-treated cells. B) Quantification of H3K9ac levels in prophase cells treated with DMSO (n=33 cells, 4 replicates) or VPA (n=25 cells, 3 replicates for VPA; p=0.0083), to determine the efficacy of VPA treatment in increasing histone acetylation. C) Representative images of SMC2 immunostaining in controls and SMC2-depleted cells in interphase and prophase, to demonstrate the efficacy of the RNAi treatment. D) Quantification of chromosome condensation using the coefficient of variation (CV) parameter. Treatment with ICRF-193 (n=49 cells, 5 replicates, p=0.0170) or VPA (n=48 cells, 5 replicates, p=0.0367) significantly decreases the CV (p<0.0001) when compared to control cells (n=52 cells, 4 replicates), reflecting an impairment in chromosome condensation. E) Quantification of chromosome condensation after depletion of SMC2 by RNAi. The CV is significantly decreased in depleted cells (n=42 cells, 5 replicates; p=0.0104) when compared to control cells (n=91 cells, 8 replicates). F) FLIM measurement of histone H4-GFP in interphase or prophase cells treated with ICRF-193 (n=20 cells for interphase and n=13 cells for prophase). Decreased fluorescence lifetime corresponds to increased chromosome compaction. G) Median of the majorant of frequency-dependent NE membrane displacements, uf, obtained by finding the maximum amplitude of the spatial Fourier transform (FT) for each frequency for DMSO (n=27 cells), ICRF-193 (n=30 cells) or VPA-treated cells (n=22 cells; p<0.0001). This FT curve shows the maximum displacement amplitude of each wavelength.

**Figure EV3 – Chromosome condensation sets the timing of mitotic entry**

A) Representative frames from a movie of a HeLa cells expressing cyclin B1-Venus/H2B-RFP during mitotic entry. B) Representative frames from movie of a HeLa cell expressing DHC-GFP and stained with SiR-DNA during mitotic entry. Note the accumulation of dynein on the NE (yellow arrowheads). C) Kymograph of the cell in B), highlighting the accumulation of dynein on the NE prior to NEP (asterisks), as well as its accumulation on kinetochores and cell cortex after mitotic entry. Horizontal scale bar, 200 sec. Vertical scale bar, 10 μm. D) Ratio between dynein on the NE and on the cytoplasm over time in prophase cells (n=33). Note how the ratio is above 1, reflecting an enrichment on the NE at this stage. In the movies, time zero corresponds to the moment of NEP. E) Individual traces for cyclin B1 nuclear translocation for cells treated with DMSO, ICRF-193 or VPA. Time zero corresponds to the time of CDK1 inhibitor (CDK1i) washout. Values were normalized and scaled between 0 and 1, corresponding to the minimum and maximum values, respectively. F) Half-times for cyclin B1 translocation were calculated from the curve fits for DMSO, ICRF-193 (p=0.0036) and VPA (p=0.0001). G) Representative frames from movies of RPE-1 cells expressing cyclin B1-eYPF after synchronization with a CDK1 inhibitor. Following washout, cells were filmed during mitotic entry. Time lapse is 5 minutes. Time is in min and scale bars are 100 μm. H) Quantification of the time from CDK1i washout to cyclin B1 nuclear translocation in RPE-1 cells (n=232 cells, 5 replicates for DMSO; n=149 cells, 3 replicates for VPA; p=0.0045). I) Representative frames from movies of cells expressing H2B-GFP/tubulin-RFP that were synchronized in late G2 using CDK1i and incubated with DMSO, ICRF-193 or VPA overnight. Following CDK1i washout, cells progressed to prophase, in medium still containing either DMSO, ICRF-193 or VPA. Time zero corresponds to washout of CDK1i. J) Quantification of the time from CDK1i washout to NEP for DMSO (n=70 cells, 9 replicates), ICRF-193- (n=84 cells, 3 replicates, p=0.0021) or VPA-treated cells (n=85 cells, 4 replicates, p=0.0038). NEP is determined as the moment soluble tubulin enters the nuclear space, reflecting permeabilization of the NE. K) Cumulative percentage of cells treated with DMSO, ICRF-193 or VPA that enter mitosis (NEP) after CDK1i washout. L) Representative frames of HeLa cells expressing DHC-GFP cells during mitotic entry, after treatment with DMSO (top panel) or ICRF-193 (bottom panel). Yellow arrowheads indicate the NE. Right panels show kymographs highlighting the absence of dynein on the NE of ICRF-193 treated cells. Horizontal scale bar, 10 μm. Vertical scale bar, 200 sec. Representative frames from movies of cells expressing DHC-GFP and stained with SiR-DNA from controls (M) and ICRF-193 (N) treated cells. Note how dynein still accumulates at kinetochores (yellow arrowhead) and cell cortex (red arrowhead) after ICRF-193 treatment.

**Figure EV4 – Tension on the NE requires SUN proteins**

A) Western blotting analysis of Lamin A levels following depletion by RNAi. B) Representative images of nuclei from cells depleted of Lamin A. C) Representative examples of inwards (pink vectors) or outwards (red vectors) NE membrane displacements of a control cell (top) and a cell depleted of Lamin A (bottom). Right panels show a map of the displacement amplitude, u, obtained for all time frames. The y axis corresponds to the arc around the median NE membrane. D) Median of the majorant of frequency-dependent displacements, uf, obtained by finding the maximum amplitude of the spatial Fourier transform (FT) for each frequency for controls (n=24) and Lamin A depleted cells (n=21; p<0.0001). This FT curve shows the maximum displacement amplitude of each wavelength. E) Representative immunofluorescence images of the expression of KASH-ΔL (left panel) and DN-KASH (right panel) constructs in HeLa cells and their respective impact on Nesprin-2 localization on the NE. F) Representative immunofluorescence images of cPLA2 levels on the nucleus in cells expressing KASH-ΔL or DN-KASH. G) Quantification of cPLA2 fluorescence intensity on the NE and nucleus for cells expressing KASH-ΔL or DN-KASH (n=46 cells, 3 replicates for KASH-DL; n=42 cells, 3 replicates for DN-KASH, p=0.0174 for NE and p=0.0288 for nucleus). H) Representative immunofluorescence images of cPLA2 levels on the nucleus following SUN2 RNAi. I) Quantification of cPLA2 fluorescence intensity on the NE and nucleus for controls (n=41 cells, 3 replicates) and SUN2 depleted cells (n=33 cells, 3 replicates; p=0.0083 for NE and p=0.0495 for nucleus). Scale bars, 10 μm.

**Figure EV5 – SUN proteins regulate dynein loading through the Nup133-CENP-F pathway**

A) Representative immunofluorescence images of SUN1, Lamin A and DAPI for controls (left panel) and SUN1 depleted cells (right panel). B) Quantification of SUN1 levels on the NE of controls (n=41 cells, 4 replicates) and SUN1 RNAi cells (n=46 cells, 4 replicates, p<0.0001). C) Quantification of SUN2 levels on the NE of controls (n=27 cells, 3 replicates) and SUN2 RNAi cells (n=30 cells, 3 replicates; p<0.0001). D) Representative immunofluorescence images showing dynein and SUN1 localization on the NE for control cells and SUN1-depleted cells. E) Analysis of chromosome condensation using the coefficient of variation (CV) for controls, (n=104 cells, 8 replicates), SUN1 RNAi (n=40 cells, 5 replicates; p=0.0216) and SUN2 RNAi (n=60 cells, 4 replicates; p=0.0414). Depletion of SUN proteins leads to a small but significant increase in CV, reflecting increased chromosome condensation. F) Quantification of dynein fluorescence intensity on the NE for controls (n=32 cells, 5 replicates) and SUN1/SUN2 shRNA without (n=18 cells, 3 replicates; p=0.0034) or with confinement (n=15 cells, 3 replicates; p=0.0171). Note how confinement does not restore dynein on the NE after SUN1/SUN2 are depleted. G) Quantification of the percentage of prophase cells showing NE or nucleoplasmic localization of CENP-F for controls, ICRF-193- or VPA-treated cells. H) Immunofluorescence analysis of NudE/EL localization on the NE after treatment with ICRF-193. In control prophase cells, NudE/EL localize to the NE (yellow arrowhead). This localization is lost after treatment with ICRF-193 (right panels). I) Quantification of NudE/EL fluorescence intensity on the NE after treatment with ICRF-193 (n=20 cells, 4 replicates; p=0.0416), in comparison with control cells (n=16 cells, 3 replicates for controls). J) Quantification of the coefficient of variation (CV) for controls (DMSO, n=32 cells, 3 replicates), ICRF-193 (n=38 cells, 3 replicates, p=0.0435) or VPA (n=35 cells, 3 replicates, p=0.0269). K) Quantification of the percentage of prophase cells showing NE or nucleoplasmic localization of CENP-F in control cells, SUN1-depleted or SUN2-depleted cells. L) Representative western blotting analysis of BicD2 levels in controls and BicD2-depleted cells. M) Representative western blotting analysis of NudE/EL levels in controls and NudE/EL-depleted cells.

## Legend for Movie S1 – Chromosome condensation controls the nuclear translocation of cyclin B1

Representative movie of HeLa cells expressing cyclin B1-Venus after treatment with DMSO (control), ICRF-193 or VPA. Cells were previously synchronized in G2 using a CDK1 inhibitor (Ro-3306). Movie starts immediately after inhibitor washout (time zero). Time lapse is in min and scale bar is 10 μm. Note the severe delay in cyclin B1 translocation after treatment with either ICRF-193 or VPA.

