## Supplementary figures and images for "Chromosome condensation mechanically primes the nucleus for mitosis"

### Supplementary Figures

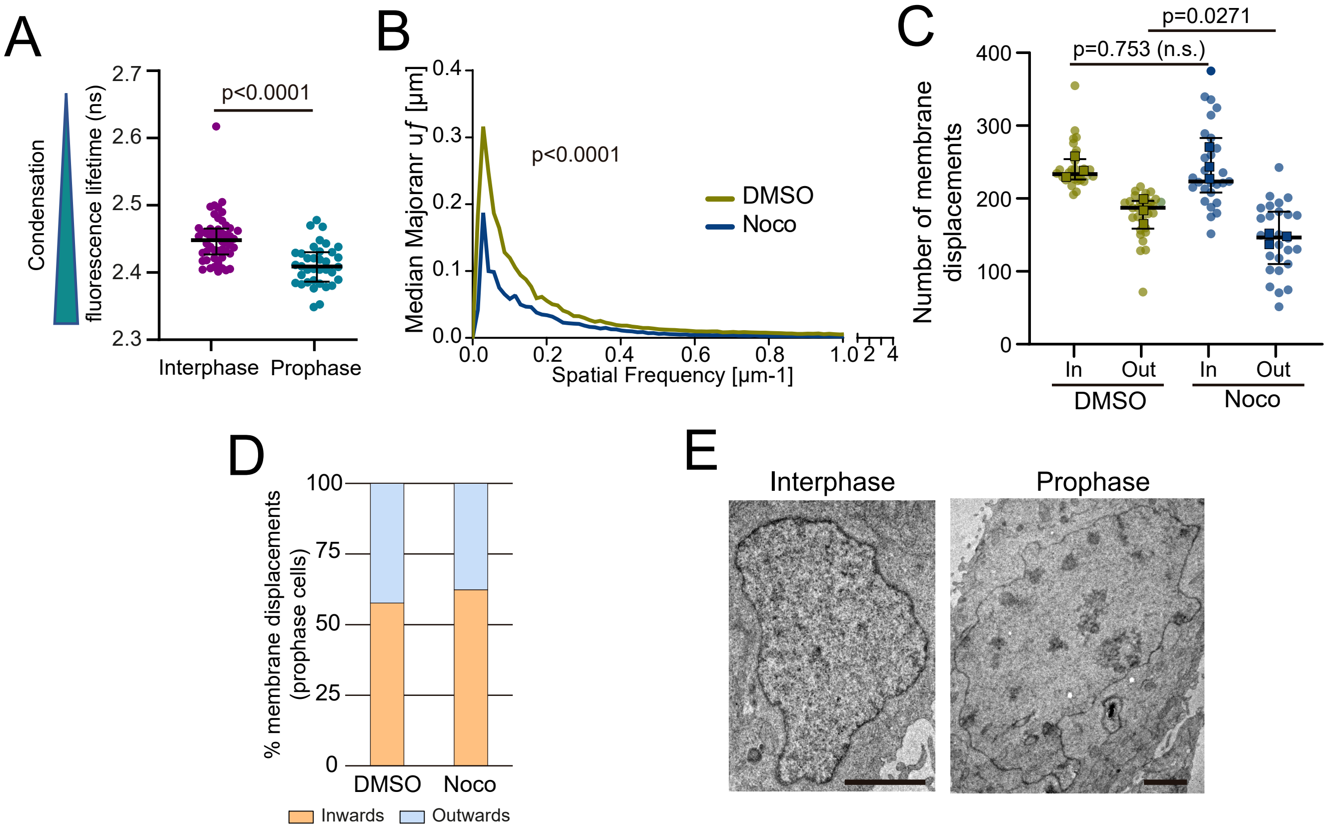

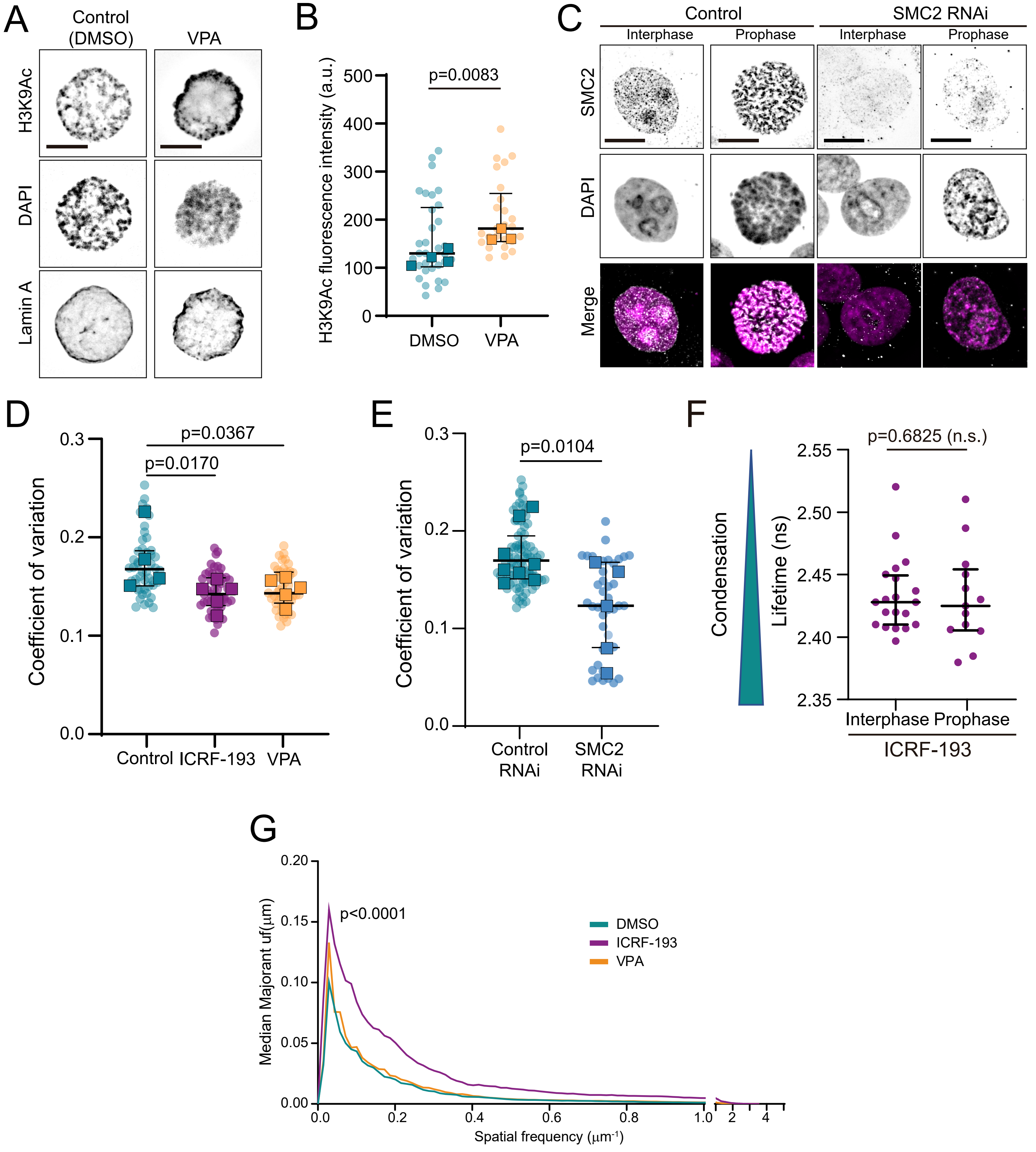

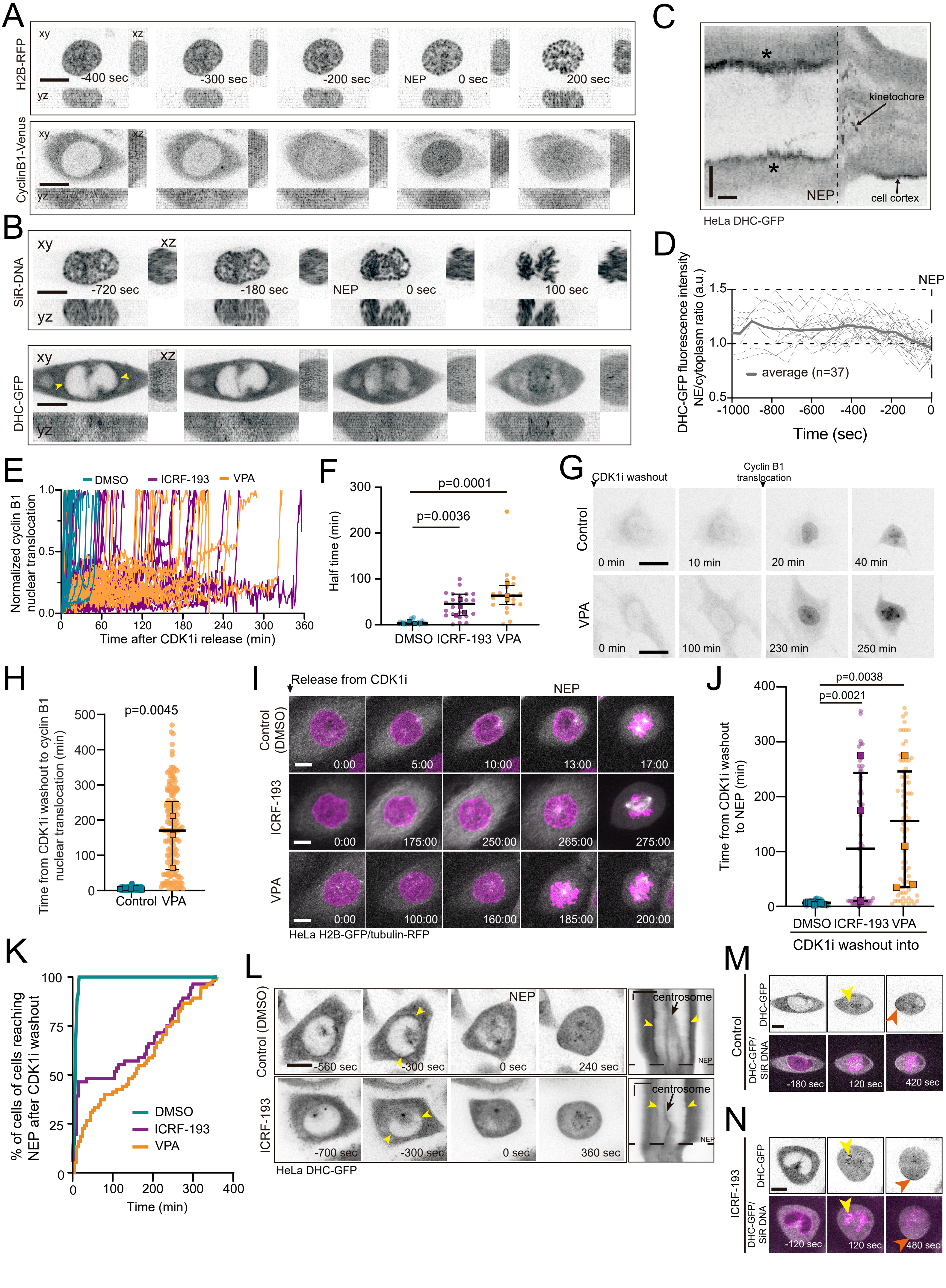

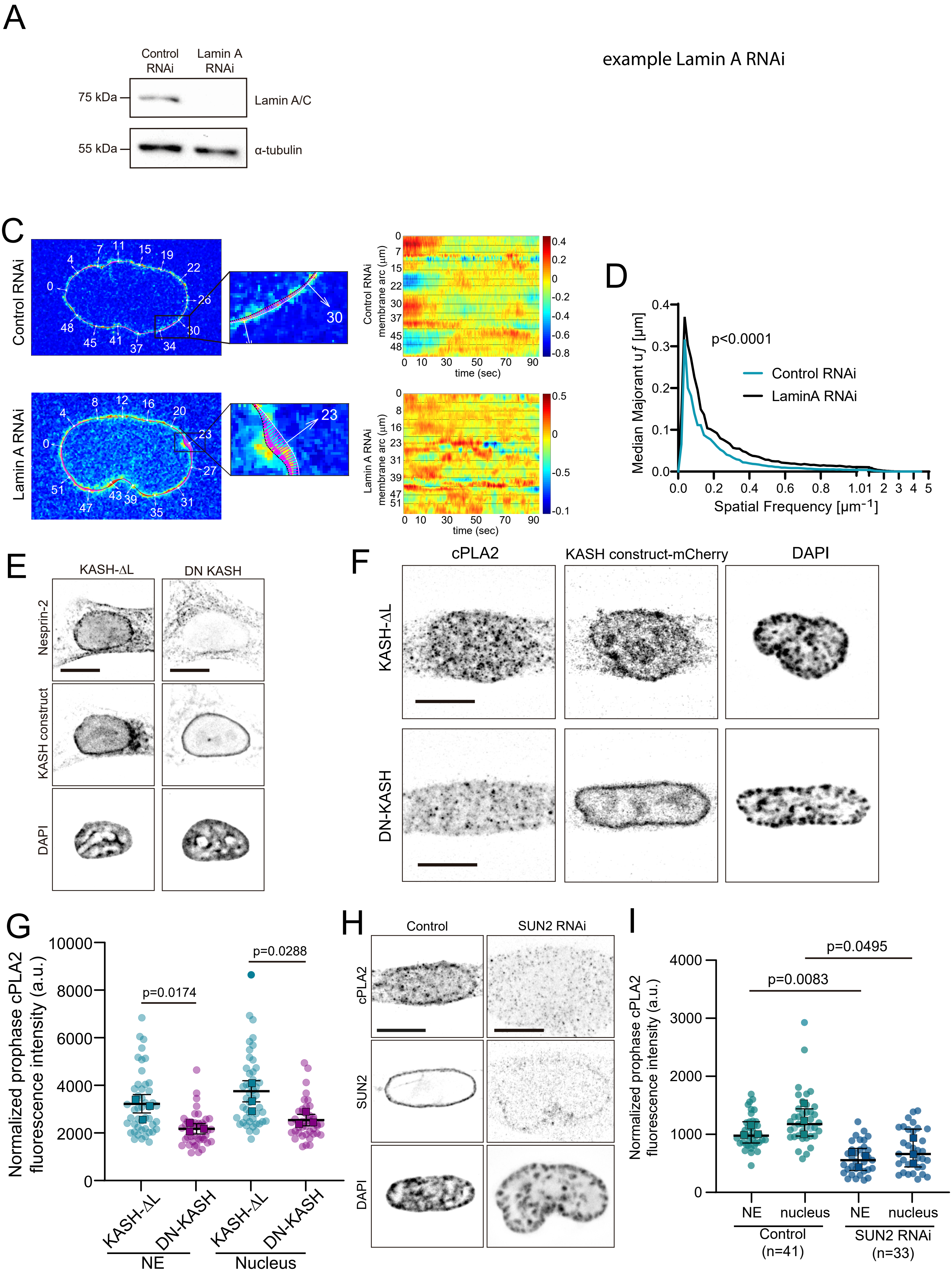

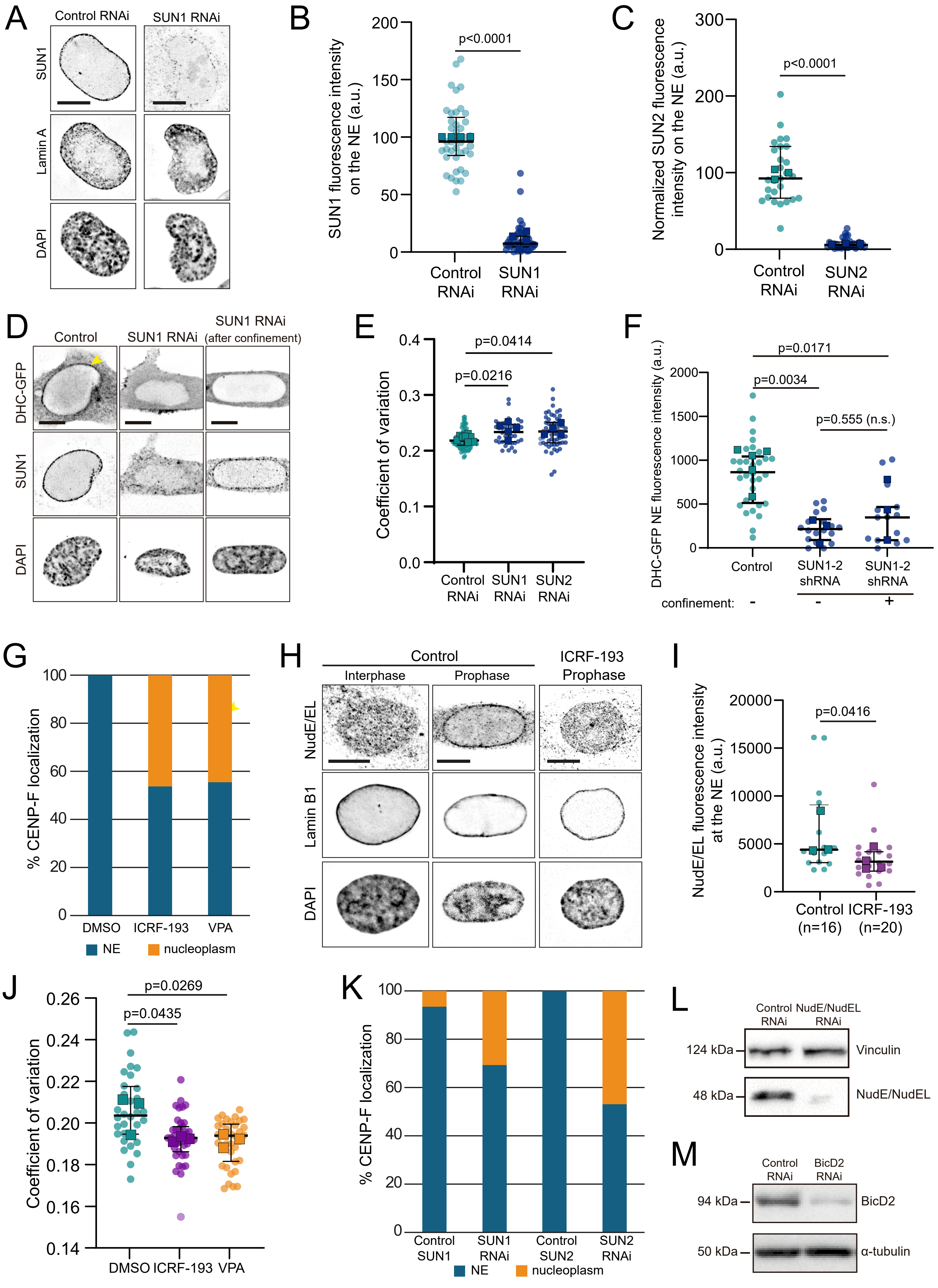
